# Domestication reshaped induced defense signaling in the apple-aphid interaction

**DOI:** 10.64898/2026.09.01.748548

**Authors:** Ronan Dadole, Anthony Venon, Xilong Chen, Maxime Criado, Amandine Hansart, Sergio Olvera-Vazquez, Yonghua Jiang, Mike Anaya, Anush Nersesyan, Anamaria Roman, Tudor-Mihai Ursu, Enrique Dapena, Natalia Conde e Silva, Amandine Cornille

**Affiliations:** Université Paris-Saclay, INRAE, CNRS, AgroParisTech, GQE - Le Moulon, 91190 Gif-sur-Yvette, France; Division of Science, New York University Abu Dhabi, Saadiyat Island, Abu Dhabi, United Arab Emirates; Department of Conservation of Genetic Resources of Armenian Flora, A. Takhtajan Institute of Botany, Acharyan Street 1, 0063 Yerevan, Armenia; Institute of Biological Research Cluj-Napoca, Branch of National Institute of Research and Development for Biological Sciences RO, 48 Republicii Street, 400015 Cluj-Napoca, Romania; Servicio Regional de Investigación y Desarrollo Agroalimentario, Ctra. AS-267, PK 19, Villaviciosa, 33300 Asturias, Spain

**Keywords:** domestication, aphid resistance, induced defense signaling, coevolution, transcriptomics, perennial crops

## Abstract

- Domestication can shift the balance between constitutive and inducible plant defense, but how this shift is reflected in defense-signaling gene expression remains poorly understood in perennial crops. We investigated how domestication altered induced transcriptional defense responses in apple to its major pest, the rosy apple aphid (*Dysaphis plantaginea*).
- We paired wild (*Malus orientalis*, and two geographically differentiated *M. sylvestris* populations from France and Romania) and cultivated (cider and dessert *M. domestica*) apple genotypes with two *Dysaphis plantaginea* genotypes collected from the same French and Romanian regions as the wild *M. sylvestris* populations sampled, a reciprocal design that is logistically demanding in a long-lived perennial host and its clonally propagated aphid pest, combining controlled infestation experiments with phenotypic measurements, paired host-aphid RNA-seq, co-expression network analysis, and genome-wide selection scans.
- Aphid fitness and host resistance varied strongly across host populations: domesticated apples supported higher aphid performance, whereas Romanian *M. sylvestris* showed the highest resistance, revealing marked host–aphid genotype-specific asymmetric compatibility. Cultivated apples mounted extensive inducible transcriptional reprogramming of hormone-signaling and secondary-metabolism pathways, whereas wild populations showed weaker transcriptional perturbation despite greater resistance, consistent with a shift toward more constitutive defense in the wild. Cross-species co-expression analyses identified coordinated host-aphid modules linking host defense-signaling genes, including a WRKY70 homolog and a CNL-class resistance gene, to aphid transcripts with effector-like annotations.
- Together, these results indicate that domestication shifted the balance between constitutive and inducible defense signaling in apple, offering a mechanistic entry point into how induced immunity to phloem-feeding insects is reshaped in a long-lived, clonally propagated perennial crop.

## Introduction

Domestication is a major evolutionary process during which humans select plants for agronomic traits such as yield, fruit size, taste, or productivity (Gaut *et al*., 2015). This process frequently modifies interactions between plants and their biotic environment, including pathogens and herbivores (Herms and Mattson, 1992; Turcotte *et al*., 2014). One long-standing hypothesis predicts that domestication may reduce resistance to parasites because artificial selection favors growth and productivity at the expense of defense (Herms and Mattson, 1992). However, empirical studies have shown that the consequences of domestication for resistance vary widely among crops and parasite systems (Turcotte *et al*., 2014). Most current knowledge regarding domestication and defense evolution originates from annual crops, yet perennial fruit trees differ substantially from annual species in their life-history traits and domestication trajectories (Miller and Gross, 2011). Perennial crops are characterized by long juvenile phases, frequent clonal propagation, high levels of introgression with wild relatives, and extensive standing genetic diversity (Miller and Gross, 2011). The evolutionary consequences of domestication in perennial crops may therefore fundamentally differ from those observed in annual species. Meta-analytical evidence suggests that while domestication generally reduces herbivore resistance across crops, the magnitude of these effects varies among life-history categories, with woody perennials potentially experiencing weaker or more context-dependent changes compared to herbaceous annuals (Turcotte *et al*., 2014; Whitehead *et al*., 2017).

Plant defenses against herbivores and pathogens can broadly be divided into constitutive (or basal) defenses, which are present before attack and provide a pre-existing level of protection, and inducible defenses, which are activated or enhanced following perception of an attacker. Constitutive defenses may involve structural barriers, defensive metabolites, or basal expression of defense-associated genes, whereas inducible defenses rely on attack-triggered signaling and transcriptional reprogramming. Inducible responses involve complex regulatory networks, including pattern-triggered immunity (PTI), wound signaling pathways, hormone-mediated defense responses, and effector-triggered immunity (ETI) (Miller *et al*., 2017). Activation of these pathways induces coordinated changes in the expression of genes associated with signaling, secondary metabolism, stress responses, and resistance functions. In particular, genes involved in jasmonic acid, salicylic acid, and abscisic acid signaling pathways, reactive oxygen species (ROS) detoxification, phenylpropanoid biosynthesis, and receptor-like kinase signaling are frequently mobilized during plant defense responses. In parallel, resistance-associated genes, notably nucleotide-binding leucine-rich repeat (NLR) receptors, contribute to attacker recognition and the activation of downstream immune responses (Miller *et al*., 2017). The relative investment in constitutive and inducible defenses can vary among populations and species and may itself evolve in response to local biotic environments. Constitutive defenses may provide immediate protection but incur continuous metabolic costs, while inducible defenses can reduce these costs in the absence of herbivores but depend on the speed and effectiveness of their activation. Accordingly, some populations may maintain higher basal activity of defense-associated pathways, whereas others rely more strongly on inducible responses following infestation (Rasmann *et al*., 2015). Defense-associated genes can also represent important targets of natural selection, with genes involved in stress signaling, secondary metabolism, resistance pathways, and regulatory responses frequently displaying signatures of positive or balancing selection in natural plant populations. These alternative defense strategies may therefore reflect distinct evolutionary histories, ecological contexts, or trade-offs between growth and defense. Understanding how domestication has reshaped these regulatory and adaptive strategies remains a major challenge, particularly in perennial crops where long generation times, clonal propagation, and recurrent introgression with wild relatives may influence the evolution of defense-associated genes and regulatory networks (Miller and Gross, 2011; Gaut *et al*., 2015).

Aphids are major agricultural pests capable of rapidly adapting to host defenses through highly specialized interactions with plants (Miñarro and Dapena, 2007; Olvera-Vazquez *et al*., 2021). As phloem-feeding herbivores, aphids establish intimate interactions with host tissues and can counteract plant immune responses by secreting salivary effectors that manipulate host signaling and facilitate feeding (Miller *et al*., 2017). These interactions can generate reciprocal selective pressures between host and parasite genomes. Host populations may evolve different defense strategies, whereas aphid populations may undergo adaptive specialization to particular host species or genotypes. Integrating host and parasite genomic and transcriptomic responses therefore provides a framework for identifying the molecular processes underlying such interactions (Multari *et al*., 2026).

The cultivated apple, *Malus domestica* Borkh., is among the most economically important fruit crops worldwide. Previous genomic studies demonstrated that cultivated apples originated from *Malus sieversii* in Central Asia and subsequently experienced extensive introgression from multiple wild relatives, notably *M. orientalis* and *M. sylvestris*, during westward diffusion along the Silk Road (Cornille *et al*., 2012, 2015, 2019; Chen *et al*., 2023, 2026). The rosy apple aphid, *Dysaphis plantaginea*, is one of the most damaging pests of cultivated apple production (Miñarro and Dapena, 2007; Olvera-Vazquez *et al*., 2021). Recent demographic analyses revealed a complex evolutionary history for this aphid, with European populations representing the ancestral lineage, followed by divergence between Mediterranean and Middle Eastern populations, and a recent expansion associated with the diffusion of the cultivated apple. These demographic patterns suggest that the spread of domesticated apples may have created new ecological opportunities that favored aphid divergence and adaptive specialization (Olvera-Vazquez *et al*., 2021).

Testing this directly favors a reciprocal design that pairs wild and cultivated host populations with parasite genotypes of matched geographic origin. Such designs remain rare in plant-insect systems and are particularly difficult to implement in long-lived perennial trees and their aphid pests. Identifying appropriate matched wild source populations first required prior population-genomic characterization of *M. sylvestris* and *M. orientalis* (Cornille *et al*., 2012, 2015, 2019), and establishing comparable wild and cultivated plant material required a multi-month stratification and propagation pipeline standardized across seed-derived wild seedlings and vegetatively propagated cultivars. Because *D. plantaginea* reproduces parthenogenetically and field populations comprise mixtures of genotypes, testing matched host-aphid pairs further required establishing single-clone aphid lineages from live colonies collected in France and Romania, molecularly validating their species identity, and synchronizing their development so that infestation could be applied simultaneously across all host populations. Combining phenotypic, transcriptomic, and population-genomic approaches within this reciprocal design offers an opportunity to test for signatures of local host-parasite coadaptation that has rarely been achieved in a perennial crop-pest system.

Here, we investigated how domestication altered apple responses to aphid infestation by integrating phenotypic analyses, transcriptomics, co-expression networks, and selection signatures. Specifically, we asked: Does domestication alter baseline defense-associated expression and inducible transcriptional responses? Do wild and cultivated apples differ in resistance and phenotypic responses to aphid infestation? Are host-aphid interactions associated with coordinated co-expression modules? Do candidate host and aphid interaction genes exhibit signatures of selection?

We show that domestication strongly reshaped apple defense-associated regulatory responses and altered compatibility with aphid populations. Cultivated apples showed extensive inducible transcriptional reprogramming and higher aphid susceptibility, whereas wild populations showed reduced aphid fitness and weaker transcriptomic perturbation, consistent with more constitutive or efficient defense strategies. In parallel, integrated host-aphid co-expression analyses and selection scans identified candidate host-aphid interaction modules linking host defense genes and aphid genes under adaptive evolution.

## Material and methods

### Plant material and growth

Following a three-month stratification, wild apple seeds were sown in Jiffy pellets (Jiffy Products, Norway) within 20-hole arrays. The study evaluated 104 plants across five populations: 54 wild seedlings (*Malus orientalis*, Romanian *M. sylvestris*, and French *M. sylvestris*) grown directly from seed, and 50 *M. domestica* cultivars (18 cider and 32 dessert cultivars) grafted onto M9 Pajam rootstocks (Table S1, Fig. 1). Wild populations were represented by two to three maternal families, with individual seedlings derived from seeds collected from the same mother tree, whereas cultivated populations were represented by clonally propagated cultivars. Multiple plants from each maternal family or cultivar were distributed across experimental conditions. Grafting was deliberately retained for cultivated genotypes to mirror real-world agronomic/domestication practices, whereas wild populations remained own-rooted to reflect natural growth conditions.

**Fig. 1.**
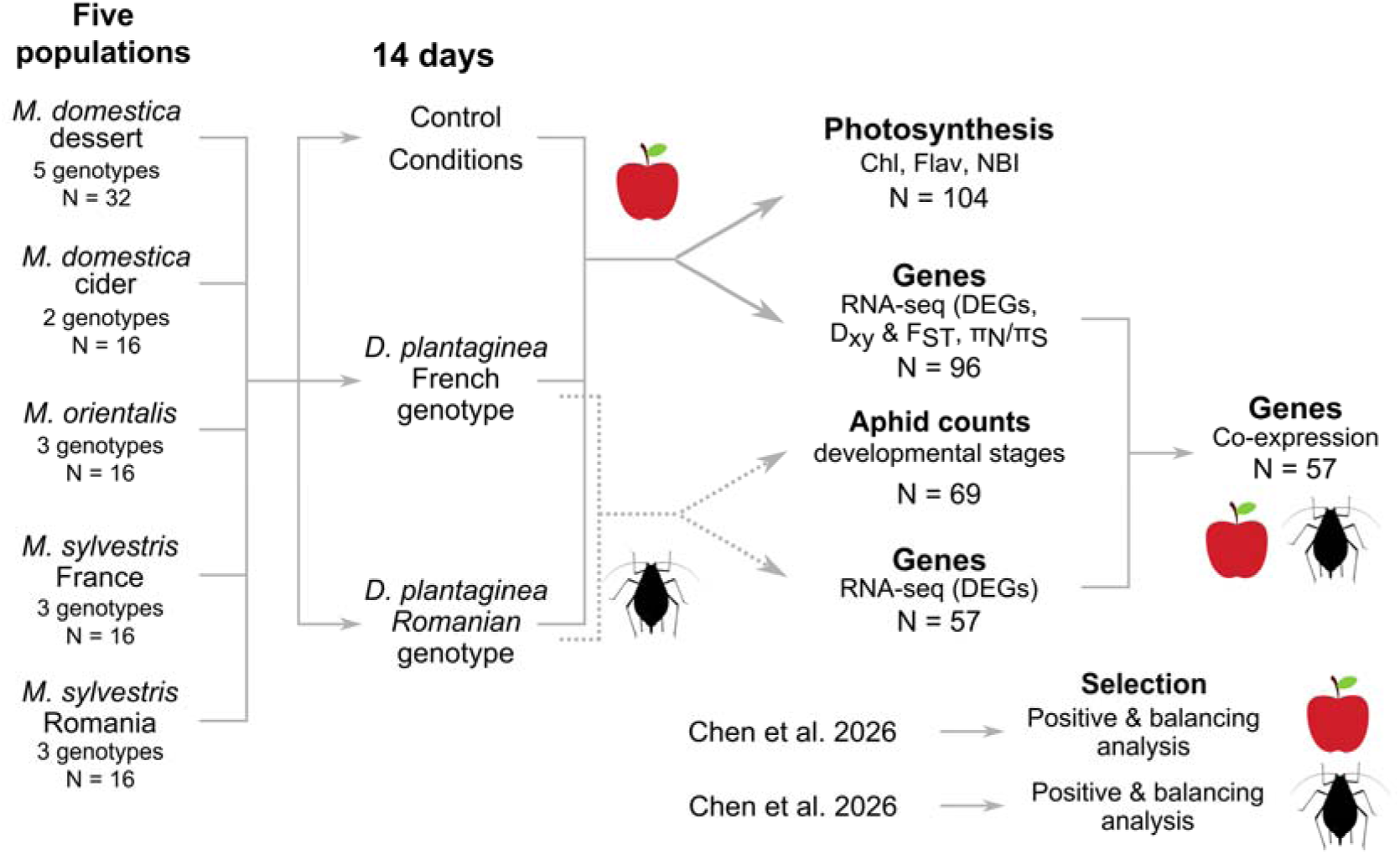
Experimental design used for infestation assays and transcriptomic analyses. Wild and cultivated apple populations were exposed to either French or Romanian aphid genotypes or maintained under non-infested control conditions. Leaf phenotypes, aphid abundance, transcriptomes, and co-expression patterns were subsequently analyzed. Photosynthetic traits included Leaf Chlorophyll content (Chl), Flavonoid content (Flav), and nitrogen balance index (NBI). Alt Text: Experimental workflow diagram illustrating five apple populations exposed for 14 days to control conditions or infestation by one of two *Dysaphis plantaginea* aphid genotypes (French and Romanian), leading to downstream analyses of photosynthesis traits, aphid counts, RNA-seq gene expression, co-expression networks, and positive and balancing selection analyses from Chen et al. 2026.

All plants were grown for three months (mid-April to mid-July 2021) in a shared, climate-controlled growth chamber (22 ± 1 °C, 60 ± 5% RH, 16:8 h L:D photoperiod, and 40–60 µmol m⁻² s⁻¹ light intensity). To minimize micro-environmental variation, individual arrays were rotated daily, and plants were watered weekly.

### Aphid rearing and validation

In April 2021, wild colonies of *D. plantaginea* were collected from Massy, France (representing the Western genetic cluster) and Cluj-Napoca, Romania (representing the Eastern genetic cluster). Colonies were initially established on *M. domestica* ‘Golden Delicious’ scions to produce fundatrigenae. To create single-clone lineages, a single female from each geographic origin was transferred to a fresh scion. Clonal lineages were maintained parthenogenetically on ‘Golden Delicious’ scions under controlled conditions: 16 h day (22 °C, 80% RH) with daylight fluorescent tubes and 8 h night (18 °C, 90% RH). Under these conditions, mature parthenogenetic females developed within 10–12 days. Aphids were handled exclusively with a fine-tipped paintbrush to prevent physical damage. Species identity for both clonal lines was molecularly validated using the cytochrome c oxidase subunit I (CO1) marker protocol from Olvera-Vazquez *et al*., (2021).

### Experimental setup and infestation

After three months, plants were transferred to a common experimental chamber and randomly assigned to one of three treatments (Fig. 1): non-infested control (n = 35), infested with the French *D. plantaginea* genotype (n = 35), or with the Romanian *D. plantaginea* genotype (n = 34). Infested plants were inoculated with three freshly emerged, synchronized adult aphids and maintained under the same environmental rearing conditions. Due to space and time constraints, plant replicates were split into two experimental batches separated by three days.

### Phenotypic measurements and statistical analyses

To characterize both sides of the plant–aphid interaction, we quantified aphid abundance as a measure of aphid performance, together with three plant physiological traits reflecting the plant response: chlorophyll content (Chl), flavonol content (Flav) and nitrogen balance index (NBI). Aphid colony growth was quantified on infested plants using category 5 (larvae) of the classification criteria (Fig. S1).

The superficial chlorophyll (Chl) content, superficial flavonol (Flav) content, and Nitrogen Balance Index (NBI) were measured on 10 leaves per plant, when possible, before infestation and at the sampling date (i.e., 14 days post-infestation). These traits were measured using a portable Dualex® leaf clip (Force-A, Orsay, France), which combines fluorescence signals across multiple excitation bands to quantify pigments and has been previously calibrated for apple trees (Hamann *et al*., 2018). The superficial chlorophyll content represents the chlorophyll concentration in the leaf epidermis (µg/cm²), and the superficial flavonol content is an index of the flavonoid concentration (µg/cm²) in this upper layer, which is related to phenol accumulation and UV protection. The NBI, used as a proxy for estimating foliar nitrogen content, was calculated as the ratio of chlorophyll to flavonols: NBI = Chl/Flav. Leaf chlorophyll, flavonol content, and NBI are parameters correlated with plant carbon uptake via photosynthesis and are therefore important for assessing the stress response. Flavonol is a phenolic compound that contributes to plant vigor, acclimation, and adaptation to biotic and abiotic stress. Measurements were performed on 10 leaves, when possible, preferentially selecting leaves near the terminal shoots, and then averaged per plant. Aphids were collected on each sampling day, counted manually, and categorized by life stage according to the visual classification criteria (Fig. S1).

To test for different responses of the crop and wild populations to the different conditions, we fitted each of the three traits with the following linear mixed model (LMM) using the lme4 R package (Bates *et al*., 2015):

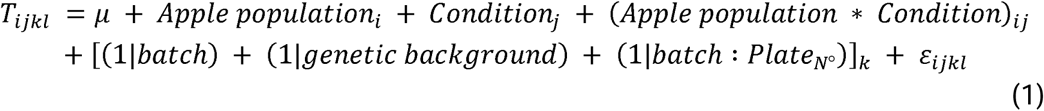

where *T_ijkl_* represents the trait of the individual from apple population *_i_* in infestation condition *_j_*, while accounting for the random effects of the individual’s batch, the genetic background (maternal family for wild seedlings and cultivar identity for cultivated apples), and the plate within the batch (batch: PlateN°). The models were fitted using a Gaussian distribution with the lme4 package in R. In cases where the interactions were statistically significant, the population and condition effects were investigated and tested separately.

We also tested for differences in aphid colony growth among cultivated and wild apple populations in response to the two aphid genotypes. We fitted a generalized linear mixed model (GLMM) of the Poisson family using the lme4 R package (Bates *et al*., 2015) as follows:

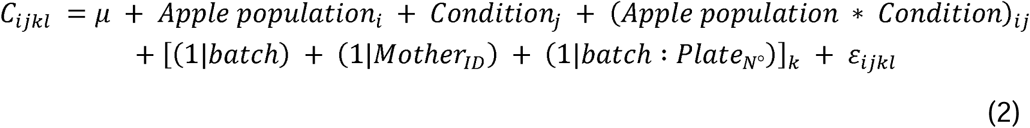

where C_ijkl_ represents the aphid count on the individual from apple population _i_ under condition _j_, while accounting for the same random effects structure as described in the plant model. Dispersion was assessed using the testDispersion function of the DHARMa R package (Hartig, 2024) (Fig. S2).

To specifically test whether the difference between aphid genotypes depended on host domestication status, we derived a binary factor domestication_status (“cultivated” for cider and dessert cultivars, “wild” for *M. orientalis* and both *M. sylvestris* populations). We then fitted a generalized linear mixed model (GLMM) of the negative binomial family using the lme4 R package (Bates *et al*., 2015) as follows:

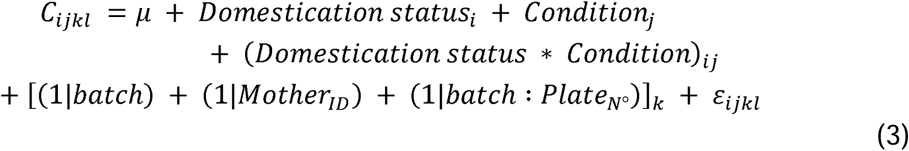

where C_ijkl_ represents the aphid count on the individual of domestication status _i_ under condition _j_, while accounting for the same random effects structure as described in the plant model.

### RNA extraction and sequencing

Leaf samples were collected after 14 days of infestation, flash-frozen in liquid nitrogen for RNA sequencing, and stored at −80 °C. In total, leaf tissue was sampled from five distinct backgrounds: cider and dessert *M. domestica* cultivars, *M. orientalis*, and both French and Romanian populations of *M. sylvestris* (Table S1). To counteract high endogenous phenolic concentrations, leaf samples were disrupted using a TissueLyser II system (Qiagen, Hilden, Germany) with the addition of polyvinylpyrrolidone (PVP40). Total RNA was extracted using the NucleoSpin RNA Plant kit (Macherey-Nagel, Düren, Germany) following the manufacturer’s protocol, incorporating an on-column DNase digestion step to eliminate genomic DNA contamination. Initial RNA quantification and purity assessments were performed using a NanoDrop spectrophotometer. Samples displaying absorbance ratios outside the threshold range of 1.8 < A_260_/A_230_ < 2.2 or A_260_/A_280_ < 1.8 were classified as poor quality and excluded from further analysis. RNA-seq libraries were prepared by Novogene with mRNA enrichment and the Novogene NGS Stranded RNA Library Prep Set (PT044 internal kit) and sequenced on a NovaSeq 6000 instrument as paired-end 150-bp reads.

### Raw RNA sequence data treatment

The quality of the raw RNA-seq reads was evaluated using FastQC (Andrews, 2010; version 0.11.7) (Table S2), and individual quality reports were consolidated using MultiQC (Ewels *et al*., 2016; version 1.11). Read filtering, adapter clipping, and quality trimming were performed using fastp (Chen, 2025; version 0.21.0); specifically, the first 9 bp of the 5’ ends were trimmed, and any reads shorter than 50 bp post-trimming were discarded. Contaminating ribosomal RNA (rRNA) sequences were identified and removed using SortMeRNA (Kopylova *et al*., 2012; version 4.3.2) against its default reference databases. High-quality non-rRNA reads were mapped to the cultivated apple reference genome (Daccord *et al*., 2017; GDDH13 version 1.1) using STAR (Dobin *et al*., 2013), and transcript abundances were quantified using featureCounts (Liao *et al*., 2014; version 2.0.3) in fragment-counting mode.

### Apple differential gene expression analyses

To evaluate the impacts of seedling population origin (comprising two cultivated and three wild populations) and experimental infestation conditions (Control, French aphid genotype, or Romanian aphid genotype) on global transcriptional output, gene expression (GE) was modeled using a general linear model:

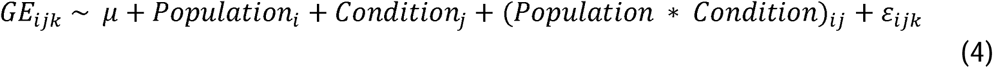

where GE_ijk_ is the number of reads associated with a gene for an individual from population _i_ under condition _j_, *μ* is the overall mean, Population _i_ is the fixed effect of the i-th population, Condition _j_ is the fixed effect of the j-th treatment condition, and *ε*_ijk_ represents the residual error. The interaction term (Population * Condition)_ij_ evaluates differential genetic architectures across populations or varying transcriptional plasticity in response to specific aphid genotypes. Differential gene expression analysis was conducted using the DiCoExpress pipeline (Lambert *et al*., 2020). Low-expressed features were filtered out using the NbCondition criterion, retaining genes with a minimum count-per-million (CPM) threshold of 5. Read counts were normalized using the Trimmed Mean of M-values (TMM) method. Statistical significance for the fixed factors (population, condition, and their interaction) was evaluated using an adjusted significance threshold (FDR = 0.05). Following normalization and filtration steps, 24,413 of the 52,741 total genomic features were retained for downstream differential expression analysis. Differentially expressed genes (DEGs) were further restricted to those exhibiting an absolute fold-change >= 2.

To confirm that processing samples across two experimental batches did not introduce confounding technical variability globally or locally, a variance partitioning analysis was conducted on the filtered matrix using the variancePartition package (Hoffman and Schadt, 2016). For each gene, normalized expression was modeled using a linear mixed-effects model:

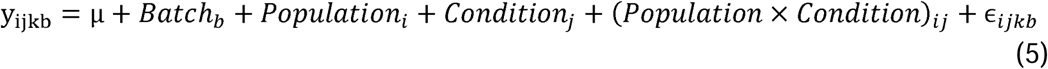

where y_ijkb_ is the log2-CPM expression value of a gene for an individual from batch_b_, population_i_, under condition_j_, μ is the baseline expression mean, Batch_b_ represents the random effect of the _b_-th batch, Population_i_ is the random effect of the _i_-th population, Condition_j_ is the random effect of the _j_-th condition, (Population x Condition)_ij_ is the random interaction effect, and ɛ_ijkb_ represents the residual error. Modeling all terms as random effects N(0, σ^2^_X_) allowed total gene expression variance to be partitioned into additive variance components:

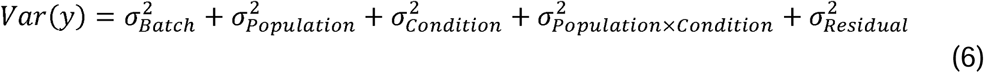

Experimental batch accounted for a negligible proportion of global expression variance across the transcriptome (mean = 1.195%, median = 0.000%) (Fig S3), confirming the absence of substantial batch effects and justifying the exclusion of batch as a covariate in the final GLM.

To assess baseline variation among the apple populations under non-stressed states, pairwise contrasts isolated the population effect exclusively within the control group. This approach yielded results that were statistically indistinguishable from those of a separate model fitted solely to control samples. To characterize the specific molecular response to herbivory, subsequent contrasts evaluated the effects of condition within each apple population independently.

### Gene ontology (GO) and function enrichment analyses

Gene Ontology (GO) term enrichment analysis was conducted using the R package clusterProfiler (Yu *et al*., 2012; version 4.6.0). Statistical significance was adjusted using the Bonferroni method, applying a strict cutoff of p < 0.05 and q < 0.05. The background gene universe was restricted to the 24,413 filtered genes evaluated in the DiCoExpress analysis. Enrichment profiles were visualized using the enrichplot package (Yu, 2025). Bubble plots were generated using the GOplot R package ((Walter *et al*., 2015); version 1.0.2), applying the reduce_overlap function with a similarity threshold of 0.75.

### Baseline and inducible gene set over-representation analysis

To test for significant overlap between baseline gene expression differences and inducible defenses, we performed over-representation analyses using one-sided Fisher’s exact tests across mutually exclusive 2x2 contingency tables. For each contrast comparing a cultivated variety (cider or dessert) against a wild population (*M. sylvestris* Romania, *M. sylvestris* France, or *M. orientalis*) in control condition, genes in the background universe (N) were cross-classified based on whether they belonged to the core set of 1,201 *M. domestica*-specific infestation-responsive DEGs, the contrast-specific baseline DEGs under control conditions, both, or neither. Tests evaluated whether core inducible genes were significantly enriched among baseline DEGs. Raw p-values were adjusted using the Benjamini–Hochberg False Discovery Rate procedure (p-adj < 0.05).

### Functional GO enrichment and response directionality

To evaluate the biological functions of the overlapping gene sets, we performed Gene Ontology (GO) enrichment analysis using the R package clusterProfiler (Yu *et al*., 2012). Enriched biological process terms were filtered at an adjusted significance threshold of p-adj < 0.05 using the Benjamini–Hochberg correction. To quantify the direction of baseline expression bias within enriched GO terms, we calculated Z-scores using the GOplot R package (Walter *et al*., 2015). The Z-score was defined as the difference between the number of genes displaying higher baseline expression in the wild population and those displaying lower baseline expression in the wild population relative to *M. domestica* under control conditions, divided by the square root of the total number of assigned genes in that term. Under this formulation, a positive Z-score indicates a higher baseline expression of the pathway in wild progenitors than in cultivated apples.

### Aphid differential expression analysis

Transcriptomic data processing for the aphid samples mirrored the workflow applied to the host plant data, with reads aligned to the *D. plantaginea* reference genome version 4.1 (BIPAA database, https://bipaa.genouest.org/sp/dysaphis_plantaginea/; Legeai *et al*., 2010; Chen, Saidi, *et al*. 2026). Raw read quality control was executed via FastQC (Andrews, 2010; version 0.11.7) and aggregated with MultiQC (Ewels *et al*., 2016; version 1.11). Quality filtering and trimming were conducted using fastp (Chen, 2025; version 0.21.0), removing 9 bp from the 5’ ends and filtering reads under 50 bp. Ribosomal RNA contaminants were systematically excluded using SortMeRNA (Kopylova *et al*., 2012; version 4.3.2) against its standard database libraries. The remaining high-quality reads were mapped to the *D. plantaginea* genome using STAR (Dobin *et al*., 2013; version 2.7.9a) in transcriptome-mapping mode (Table S2). Transcript abundance quantification from the transcriptome-aligned reads was subsequently executed using Salmon (Patro *et al*., 2017) in alignment-based mode, and gene-level and estimated number of reads were extracted. Cross-species comparisons were restricted to normalized, variance-stabilized within-gene expression profiles across matched samples and differential-expression patterns.

### Co-expression analysis

To investigate co-expression between *M. domestica* and *D. plantaginea*, a unified expression matrix was constructed by concatenating the normalized count data for both the host and the herbivore. This integrated matrix enabled simultaneous modeling of gene expression profiles across all experimental samples. Co-expression analysis was performed using the DiCoExpress pipeline, which leverages the coseq R package (Rau and Maugis-Rabusseau, 2018) to identify clusters of genes sharing similar expression patterns through Poisson mixture models.

Prior to clustering, genes were partitioned into functional subsets based on their differential expression profiles to isolate specific regulatory drivers. A custom R script automated the extraction of gene targets from the DiCoExpress directory framework, isolating unique DEGs associated with three primary experimental components. The Host Effect subset comprised genes showing differential expression across different apple population. The Treatment Effect subset included genes responsive to experimental conditions, while the Interaction Effect subset comprised genes whose treatment response was significantly modulated by the specific host-aphid combination. The normalized counts for these unique DEGs were extracted from the master matrix and used as direct inputs to the Co-expression_coseq function. This methodology enabled the identification of synchronized transcriptional shifts across the host-pathogen interface, facilitating the detection of inter-specific regulatory hubs associated with the host response, treatment impact, and their respective interactions.

### Genetic divergence between populations

To assess differentiation between populations, *F_ST_* was calculated from the mapped RNA-seq reads. The SNP dataset used was generated using the following pipeline: variants were called per sample from BAM files using GATK HaplotypeCaller v4.4.0.0 (Auwera and O’Connor, 2020) against the GDDH13 v1.1 reference genome (Daccord *et al*., 2017). Single-sample genomic VCFs (gVCFs) were generated with--emit-ref-confidence GVCF and a minimum confidence threshold of 20. Individual gVCFs were combined into a multi-sample cohort using GATK CombineGVCFs, followed by joint genotyping with GenotypeGVCFs using the -all-sites option to retain both variant and invariant positions. Raw VCFs were filtered to retain high-quality variants in a multi-step pipeline. Sites missing critical annotations (F_MISSING = 1) were removed using BCFtools v1.14 (Danecek *et al*., 2021). Remaining sites were annotated and flagged using GATK VariantFiltration based on recommended hard-filtering parameters: SNPs with QUAL < 30, QD < 2.0, ReadPosRankSum < -8.0, FS > 60.0, SOR > 3.0, or MQ < 40.0 were flagged, with InDels/mixed variants subject to adjusted thresholds (ReadPosRankSum < −20.0, FS > 200.0, SOR > 10.0). Finally, all flagged sites were removed using GATK SelectVariants, yielding a VCF containing only high-confidence variants. SNPs were then classified into synonymous and nonsynonymous variants using SnpSift (Cingolani *et al*., 2012).

*F_ST_* values among populations were computed per coding sequence, using only synonymous SNPs and invariant sites, as the default requirement of Pixy (Korunes and Samuk, 2021), to avoid bias. Weighted mean *F_ST_* values were then computed from the per-coding-sequence estimate.

### Selection analysis in apple

Genomic variant data and baseline selection signatures were derived from Chen *et al*., 2026, which provided whole-genome resequencing data for the same wild and cultivated apple accessions analyzed here. For cultivated apples, selection scan statistics were adopted directly from Chen *et al*., 2026. For the wild apple cohorts, French *Malus sylvestris*, Romanian *M. sylvestris*, and *M. orientalis*, genomic data from this same Chen *et al*., 2026 dataset were extracted, and selection analyses were re-executed using the parameters described below. Wild and cultivated selection scans therefore use the same underlying sampling scheme, ensuring full comparability between the two groups.

Signatures of positive selective sweeps in the genomes of the wild apple populations were detected using OmegaPlus (Kim and Nielsen, 2004). Before analysis, VCF files were filtered to exclude invariant sites, variants with missing data exceeding 20%, and variants with a minor allele frequency (MAF) below 0.01. OmegaPlus was applied to each chromosome individually per population using the parameters -minsnps 5, -minwin 5000, and -maxwin 100000. The grid size for each chromosome was explicitly set to partition the chromosomes into evaluation intervals of approximately 1,000 bp. Signatures of balancing selection were inferred using BetaScan (Siewert and Voight, 2017), which identifies the enrichment of variants in regions with low deviation from allele frequency and a deficit of substitutions. For this analysis, VCF files were filtered to remove invariant sites and variants with missing data exceeding 20%. To account for the background mutation rate, Watterson’s theta (θ_T_) was estimated across the genome using pixy with a sliding window size of 5,000 bp. Windows with fewer than 1,000 valid sites or undefined estimates were masked to prevent artifactual signals. VCF files were subsequently converted to the input format required by BetaScan. Standardized β scores (-std) were calculated utilizing a provided θ_T_ map, folded allele frequencies (-fold), an allele frequency core threshold of 0.15 (-m 0.15), and a window size of 1,000 bp (-w 1000).

### Identification of outlier regions and candidate genes

We assessed significance thresholds for selection analyses empirically. We considered the top 1% of the ω and β score distributions as significant outlier regions and used these thresholds for positive and balancing selection signals, respectively. To ensure data quality, pre-defined low-mappability regions, detected in Chen *et al*., 2026, were completely excluded from the selection signal results. Finally, to identify candidate genes associated with these selective sweeps, we intersected the top 1% of filtered genomic regions with the apple reference gene models. This intersection included a 500 bp upstream extension to capture associated promoter and regulatory regions.

### Positive selection analysis in aphid

Single-nucleotide polymorphisms (SNPs) for French (FR) and Romanian (RO) aphid populations were extracted from Chen, Saidi, *et al*. 2026. To assess population genetic structure, we conducted Discriminant Analysis of Principal Components (DAPC) using the R packages adegenet (Jombart and Ahmed, 2011) and vcfR (Knaus and Grünwald, 2017). Input variants were filtered from autosomal synonymous sites, excluding invariant positions (AC == 0 || AC == AN). Genetic clusters were identified using find.clusters (Jombart and Ahmed, 2011), and the number of retained principal components was optimized using optim.a.score before calculating individual cluster assignment probabilities.

Selective sweeps were evaluated across all autosomes (chr1-chr5) and the X chromosome (chrX), corresponding to *D. plantaginea* assembly scaffolds HiC_scaffold_1 through HiC_scaffold_6. Population-specific VCF files (FR and RO) were filtered using bcftools to exclude invariant sites, variants with missing data exceeding 20% (F_MISSING > 0.2), and variants with a minor allele frequency below 0.01 (MAF < 0.01). Selective sweeps were identified using OmegaPlus-M by evaluating linkage disequilibrium (LD) patterns via the ω statistic (Kim and Nielsen, 2004). Runs were executed per chromosome for each population using -minsnps 5, -minwin 5000, and -maxwin 100,000. Chromosome-specific grid sizes (43,644 for chr1; 32,832 for chr2; 30,076 for chr3; 13,988 for chr4; 13,353 for chr5; and 55,457 for chrX) were specified to achieve uniform evaluation intervals of approximately 2,000 bp.

Empirical significance thresholds were established by identifying the top 1% of ω values for each population independently (yielding empirical thresholds of ω = 11.08 for FR and ω = 7.41 for RO). Candidate sites located within low-mappability genomic regions were masked and excluded to eliminate false positives. Candidate genes associated with selective sweeps were identified by intersecting the filtered top 1% genomic regions with *D. plantaginea* reference gene models using bedtools intersect (Quinlan and Hall, 2010). Gene boundaries were extended by 500 bp upstream and downstream to capture associated promoter and regulatory sequences.

## Results

### Aphid compatibility and host resistance differed between wild and cultivated apples

We detected significant interactions between apple population and aphid genotype for chlorophyll content, flavonol accumulation, and NBI, indicating population-specific responses to infestation (Supplementary Table S3a). Per-population post-hoc comparisons revealed significant differences among experimental conditions only for the cider population and the Romanian *M. sylvestris* population (Fig. 2A; Supplementary Figs S4–S5, Table S4). Cider apples showed significant differences between control and both infested conditions for all three physiological traits, indicating a consistent physiological response to infestation irrespective of aphid genotype. By contrast, Romanian *M. sylvestris* did not differ between control and infested conditions but differed significantly between infestation by French and Romanian aphids, suggesting aphid genotype-specific responses to herbivory.

**Fig. 2.**
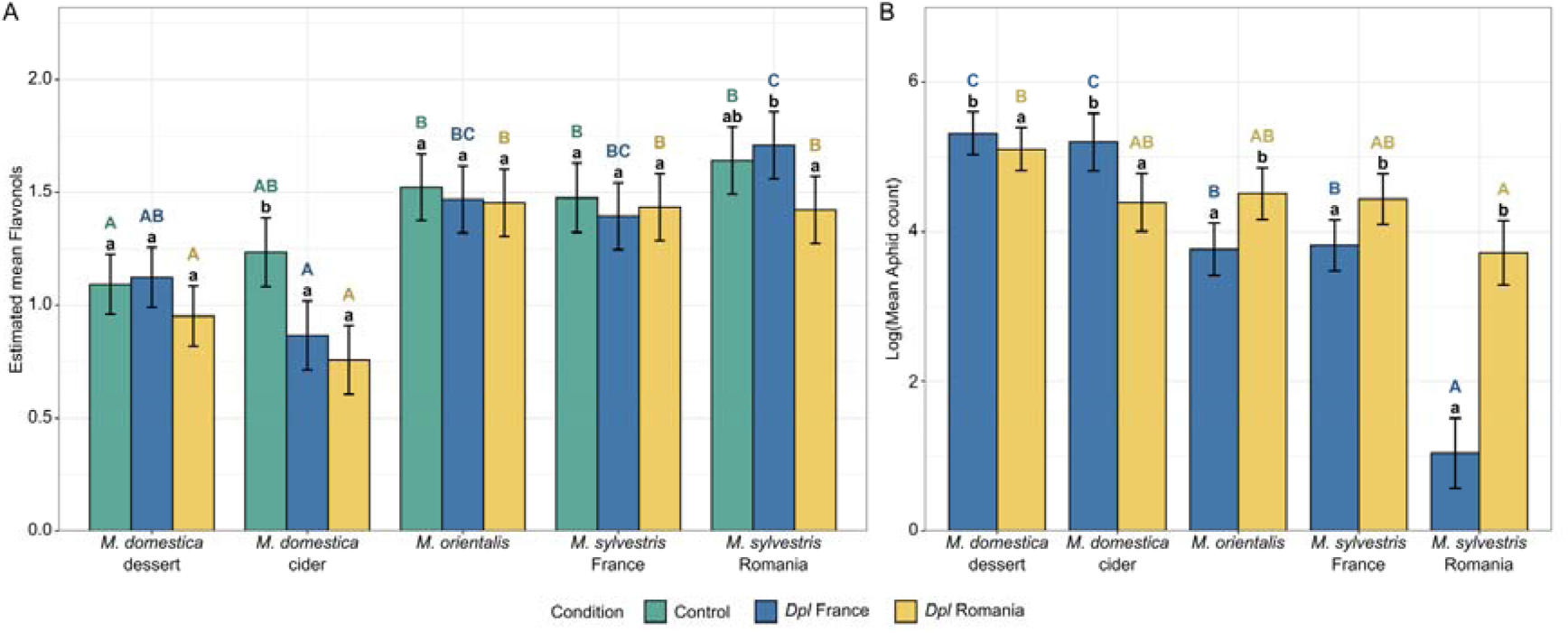
Domestication altered aphid compatibility and host physiological responses. (A) Estimated mean leaf flavonol content (± SE) and (B) estimated log mean aphid number (± SE) across five apple populations (*Malus domestica*, *M. orientalis*, and *M. sylvestris*) subjected to three conditions: uninfested control (green), infested by French aphid genotype (blue), and infested by Romanian aphid genotype (yellow). Bar heights represent the estimated marginal means (emmeans) derived from the interaction models. Error bars represent the standard error of the mean. Different letters above the bars indicate statistically significant differences based on Tukey’s HSD post-hoc tests (p < 0.05). Black lowercase letters denote significant differences among aphid treatment conditions within a single apple population. Colored uppercase letters denote significant differences across the five apple populations within a specific aphid treatment condition. Because the three physiological traits showed concordant population-specific patterns, Fig. 2A shows flavonol content as a representative response, with complete statistical results provided in Tables S3–S4. Alt Text: Two-panel bar chart showing estimated mean leaf flavonol content (Panel A) and estimated log mean aphid counts (Panel B) across five apple populations under uninfested control (green), French aphid genotype (Dpl France, blue), and Romanian aphid genotype (Dpl Romania, yellow) conditions. Bar heights represent estimated marginal means with ± SE error bars. Statistical significance (Tukey’s HSD, p < 0.05) is denoted by letters above the bars: black lowercase letters indicate significant differences among aphid treatment conditions within a single population, while colored uppercase letters indicate significant differences across populations within a specific treatment condition.

Aphid abundance analyses further demonstrated pronounced differences in host resistance (Fig. 2B). The interaction between apple population and aphid genotype was significant for aphid counts at the end of the experiment (Table S3b). Cultivated apples generally supported higher aphid abundance, whereas Romanian *M. sylvestris* exhibited the lowest aphid abundance, consistent with greater resistance (Table S5). Aphid performance depended on the host–aphid combination. The French aphid genotype reached higher abundance on cultivated than on wild apples, whereas the Romanian aphid genotype showed a weaker difference between cultivated and wild hosts and reached particularly high abundance on Romanian *M. sylvestris* relative to the French genotype (Table S5a,b). To further investigate host compatibility across domesticated and wild groups, we fitted a GLMM testing the effect of host domestication status (cultivated vs. wild) and its interaction with aphid genotype. The interaction between aphid genotype and domestication status was significant (Table S5c). Pairwise contrasts revealed that this asymmetry was primarily driven by the French aphid genotype, whose abundance differed significantly between wild and domesticated apples (estimate = 1.93, p< 0.0001), whereas no significant difference between wild and domesticated hosts was detected for the Romanian aphid genotype (estimate = 0.64, p = 0.0945) (Table S5d).

### Baseline transcriptomic divergence decouples from genomic differentiation

Principal component analyses (PCA) of normalized gene count data revealed clustering patterns that differed from population relationships previously inferred using population genomic SNP data (Cornille *et al*., 2012, 2015; Chen *et al*., 2023, 2026) (Fig. 3A). While the two *M. sylvestris* populations grouped closely together, *M. domestica* did not show an intermediary state between the wild species, with cider varieties exhibiting a more pronounced separation. Furthermore, wild *M. orientalis* samples clustered closer to *M. sylvestris* populations than to cultivated apples in PCA space.

**Fig. 3.**
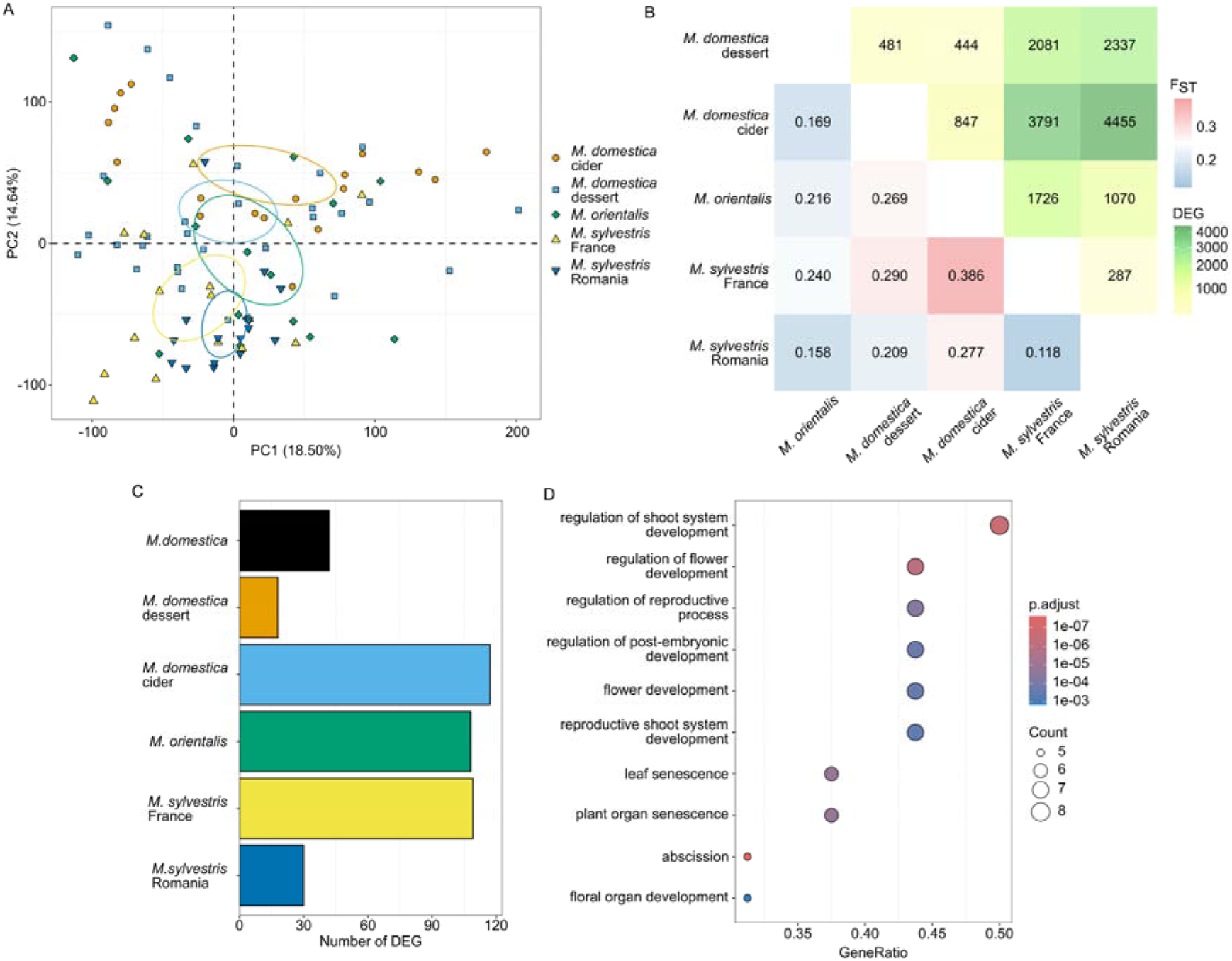
Baseline transcriptomic divergence of apples deviates from genomic divergence. (A) Principal component analysis (PCA) of gene expression profiles among wild and cultivated apple populations. (B) Heatmap of pairwise population FST (bottom triangle) and number of population-specific DEGs (top triangle) between populations. (C) Number of DEGs showing population- and *M. domestica-specific* expression levels. (D) Dot plot of Gene Ontology enrichment in DEGs showing *M. domestica-specific* expression levels. Alt Text: Four-panel figure illustrating baseline transcriptomic divergence versus genomic divergence across five apple populations (*M. domestica* dessert, *M. domestica* cider, *M. orientalis*, *M. sylvestris* France, and *M. sylvestris* Romania). Panel A shows a PCA of gene expression profiles (PC1: 18.50%, PC2: 14.64%). Panel B features a heatmap comparing pairwise genomic differentiation (F*_ST_*, bottom triangle) with differentially expressed gene (DEG) counts (top triangle) between apple populations. Panel C is a bar plot showing population and *M. domestica*-specific DEGs. Panel D shows a GO enrichment dot plot for *M. domestica*-specific DEGs.

Differential expression analyses in control conditions corroborated this transcriptomic divergence, while pairing differentially expressed gene (DEG) counts with coding sequence differentiation *F_ST_* calculated from RNA-seq SNPs revealed a decoupling between expression and sequence divergence (Fig. 3B). Across all population pairs, transcriptomic divergence showed no significant correlation with sequence differentiation (ρ = 0.297, p = 0.407). The two *M. sylvestris* populations showed the lowest transcriptomic divergence, with 287 DEGs (*F_ST_* = 0.118). In contrast, transcriptomic divergence peaked between cultivated varieties and wild *M. sylvestris* populations, specifically between the *M. domestica* cider variety and Romanian (4,455 DEGs) and French (3,791 DEGs) *M. sylvestris* populations. Dessert varieties displayed a similar, though slightly more moderate, pattern of divergence (2,337 and 2,081 DEGs, respectively). This uncoupling is illustrated by stark contrasts across population pairs: *M. orientalis* and French *M. sylvestris* showed the highest sequence differentiation (*F_ST_* = 0.386) but shared a relatively moderate number of DEGs (1,726). Conversely, *M. domestica* cider varieties and Romanian *M. sylvestris* showed low coding sequence divergence (*F_ST_* = 0.209) despite the highest transcriptomic divergence across all comparisons (4,455 DEGs).

Genes consistently differentially expressed between cultivated and wild apples were significantly enriched for developmental and reproductive functions, including floral organ development, meristem regulation, and leaf senescence (Fig. 3C, D). In particular, *AGAMOUS-like 42* homologous genes showed markedly higher expression in cultivated apples than in wild populations (Fig. S6).

### Cultivated apples trigger extensive inducible defense responses

Infestation induced transcriptomic changes across all host populations, although the magnitude of the response differed strongly among host groups (Fig. 4). PCA based on gene count showed clear separation between control and infested conditions, yet minimal separation was observed among the infested conditions themselves (Fig. 4A). Cultivated apples displayed extensive transcriptional reprogramming involving thousands of DEGs, whereas wild populations exhibited substantially weaker responses (Fig. 4B). Comparing DEG profiles in response to infestation across all populations and both aphid genotypes revealed a large overlap between *M. domestica* cider and dessert varieties, uncovering a core set of 1,201 infestation-responsive DEGs induced by both aphid genotypes (Fig. 4C). These 1,201 DEGs therefore define a shared infestation-responsive transcriptional signature detected in both cultivated apple groups and in response to both aphid genotypes.

**Fig. 4.**
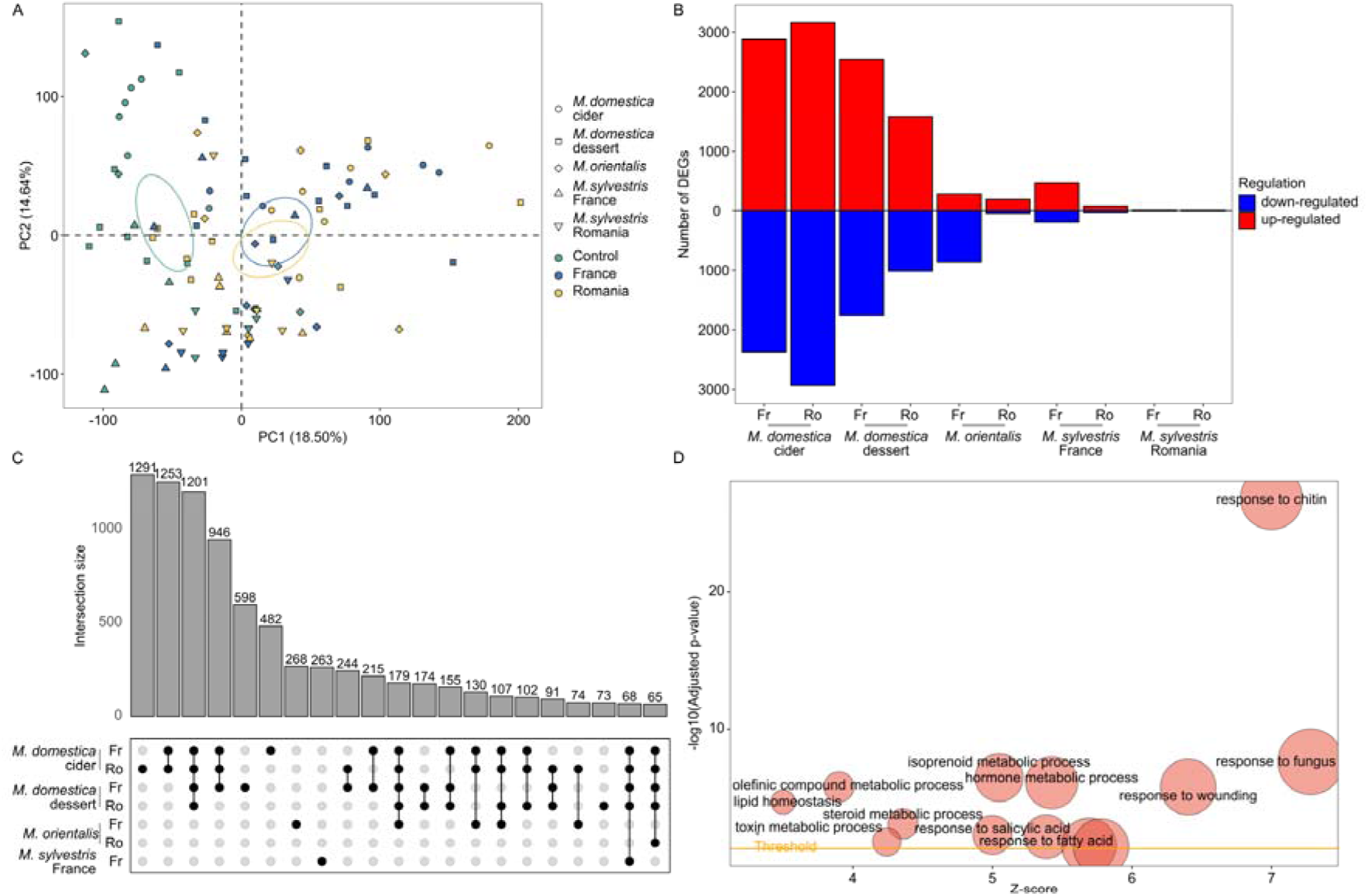
Cultivated apples exhibit extensive inducible defense reprogramming in response to aphid infestation. (A) Principal component analysis of transcriptomic responses under control and infestation conditions. Samples are primarily separated according to infestation status. (B) Number of differentially expressed genes (DEGs) identified for each population in response to infestation by French (Fr) and Romanian (Ro) aphid genotypes compared to the non-infested condition. (C) UpSet plot showing overlap among infestation-responsive DEGs. (D) Bubble plot summarizing enriched biological processes associated with induced defense responses specific to the cultivated apple. Alt Text: Four-panel figure demonstrating extensive inducible transcriptomic defense reprogramming in cultivated apples compared to wild populations upon aphid infestation. Panel A presents a PCA separating samples primarily by infestation status along PC1 (18.50%) and PC2 (14.64%). Panel B shows a bidirectional bar plot comparing up-regulated (red) and down-regulated (blue) DEG counts across five apple populations exposed to French (Fr) or Romanian (Ro) aphid genotypes. Panel C features an UpSet plot depicting the overlap among infestation-responsive DEGs between apple populations. Panel D presents a bubble plot summarizing gene ontology enrichment analysis performed infestation-responsive DEGs specific to *Malus domestica*.

Functional enrichment analyses performed using clusterProfiler on these shared DEGs revealed a strong overrepresentation of biological processes associated with biotic stress (Fig. 4D). Key enriched categories included perception and signaling mechanisms, such as response to chitin, wound signaling, and receptor-mediated signaling pathways, as well as hormone-related pathways involving jasmonic acid, salicylic acid, and abscisic acid. Z-scores indicated that genes assigned to these enriched processes were predominantly upregulated in infested individuals, consistent with activation of defense-associated pathways. We also observed enrichment for pathways involved in secondary metabolism, including terpenoids and phenolic compounds, highlighting broad metabolic reorganization during attack.

In contrast, wild populations displayed relatively few DEGs despite exhibiting lower overall aphid abundance (Fig. 2B, Fig. 4B), revealing a decoupling between transcriptional response and aphid performance. To test whether pathways induced in cultivated apples differed at baseline between wild and cultivated populations, we examined overlap between the 1,201 core inducible DEGs and DEGs between *M. domestica* varieties and each wild population under control conditions (baseline DEGs). The core set of 1,201 *M. domestica*-specific infestation-responsive DEGs was significantly over-represented among baseline DEGs identified under control conditions when compared to *M. sylvestris* Romania (cider: OR = 3.15, FDR = 2.86e-69; dessert: OR = 3.66, FDR = 1.27e-63) and *M. orientalis* (cider: OR = 4.96, FDR = 2.08e-49; dessert: OR = 2.95, FDR = 5.05e-11), whereas no enrichment was observed against French *M. sylvestris* (OR = 0.78-1.04, FDR > 0.3; Table S6). Functional GO enrichment of these overlapping gene sets identified biological processes linked to secondary metabolism, defense response, and phytohormone signaling (Fig. S7A-C), although no terms reached statistical significance for the dessert vs. *M. orientalis* contrast. Where functional enrichment was observed, positive Z-scores indicated higher baseline expression of these pathways in wild populations relative to *M. domestica*. These results indicate that some defense-associated pathways induced by infestation in cultivated apples show higher baseline expression in particular wild populations. Cider varieties additionally showed repression of photosynthesis-associated pathways during infestation (Fig. S8). Together, these analyses reveal marked differences between wild and cultivated apples in both baseline expression of defense-associated pathways and the magnitude of transcriptional responses to infestation.

### Host-aphid co-expression networks identify interaction hubs

To investigate coordinated regulatory responses between host and pest, we constructed integrated dual-transcriptome co-expression networks combining apple (*Malus* spp.) and aphid transcriptomes. Host-associated effects comprised a larger proportion of DEGs and co-expression modules than treatment-associated effects (Fig. 5A). Twenty-two modules contained both host and aphid genes, identifying cross-species gene sets with correlated expression profiles during infestation.

**Fig. 5.**
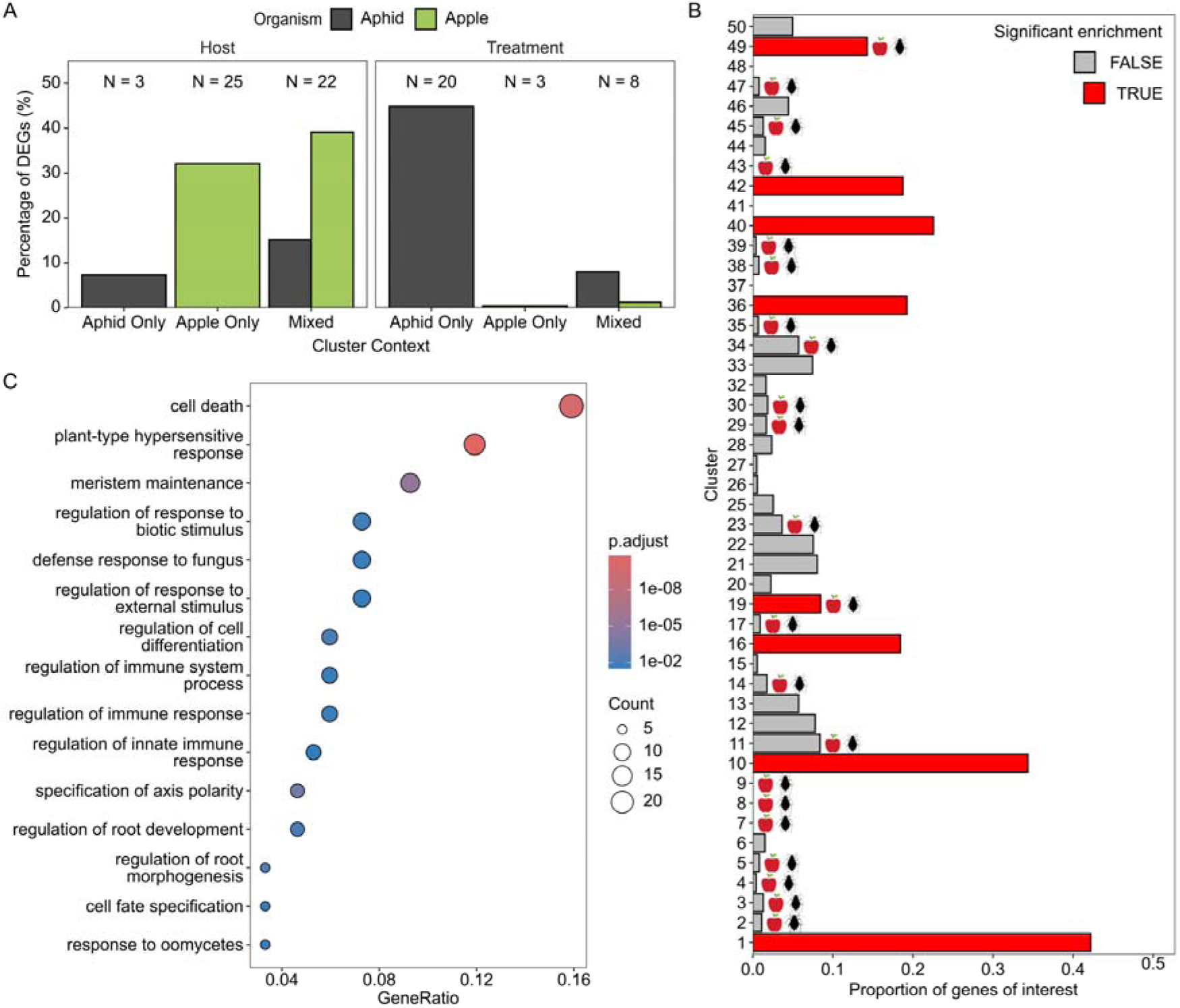
Host-aphid co-expression modules reveal candidate hubs of coevolutionary interactions. (A) Overview of co-expression modules inferred using DiCoExpress and coseq, where N corresponds to the number of modules in this context; (B) Enrichment of *M. domestica*-specific infestation-responsive genes across host-effect modules. Red modules indicate significant enrichment after FDR correction; apple and aphid symbols mark mixed co-expression modules. (C) Functional enrichment analysis of module 19 showing enrichment for defense-associated biological processes. Alt Text: Three-panel figure illustrating host-aphid co-expression modules and functional enrichment analysis. Panel A shows the percentage of DEGs across Host and Treatment cluster contexts and the number of clusters inferred using DiCoExpress and coseq. Panel B displays the proportion of *M. domestica*-specific infestation-responsive genes across co-expression clusters associated with host effect, with red bars indicating significant enrichment after FDR correction and symbols marking mixed co-expression modules. Panel C presents a dot plot of functional enrichment for module 19, highlighting defense-associated biological processes.

To test whether domestication altered host-aphid transcriptional co-regulation, we performed module enrichment analyses using the core set of 1,201 *M. domestica*-specific infestation-responsive DEGs (Fig. 5B). We detected significant enrichment in 8 of the 50 host-response modules. Notably, only two of these enriched modules comprised interspecies (mixed host-aphid) gene sets, specifically Module 19 (Table S7) and Module 49 (Table S8).

Module 19 was one of the two mixed modules significantly enriched for the 1,201 cultivated-apple infestation-responsive DEGs. This module was enriched for biological processes including hypersensitive response, programmed cell death, and symbiont-induced defense responses. Embedded aphid genes included homologs of trypsin-like proteases, fatty acyl-CoA reductases, plant cell wall-degrading enzymes, and structural proteins annotated as related to salivary sheath formation and host tissue manipulation.

To assess whether these interspecies modules overlapped with candidate regions showing signatures of balancing selection, we intersected host genome-wide scans for balancing selection with DEGs assigned to mixed host-aphid modules. This analysis identified 15 DEGs within interspecies co-expression networks (Table S9), including three candidate resistance (*R*) genes, two of which were harbored within Module 19. Among them, *MD02G1164700*, encoding a coiled-coil nucleotide-binding site leucine-rich repeat (CNL) protein, exhibited a striking host- and parasite-genotype-dependent expression profile. Specifically, *MD02G1164700* was significantly induced upon infestation in dessert varieties regardless of aphid genotype, whereas in cider varieties significant induction was detected with the French aphid genotype. In contrast, wild apple populations showed no change in gene expression following infestation.

To contrast the maintenance of balanced polymorphisms with directional evolutionary pressures, we similarly evaluated host genes under positive selection across host lineages. Intersecting positively selected genes with infestation-responsive DEGs identified more overlap in *M. domestica* (64) than in the wild populations, which harbored very few (two in *M. orientalis*, two in Romanian *M. sylvestris*, and none in French *M. sylvestris*) (Fig. 6A; Table S10). Although these positively selected DEGs yielded no overall functional enrichment and exhibited no overlap with canonical *R*-genes, three intersected directly with the core set of 1,201 infestation-responsive DEGs characteristic of cultivated apples (*MD07G1234700*, *MD01G1084700*, and *MD01G1207300*). Protein homology searches against *A. thaliana* revealed that two of these core candidates showed homology to transcriptional regulators with established roles in hormone and developmental signaling : *MD07G1234700* showed high homology to *WRKY70*, an essential defense- and senescence-related transcription factor coordinating salicylic acid and jasmonic acid signaling crosstalk (Besseau *et al*., 2012), while *MD01G1084700* shared homology with *DAG1*, a DOF-type transcription factor involved in seed germination and hypocotyl elongation through the control of abscisic acid, ethylene, and auxin signaling (Lorrai *et al*., 2018). Notably, *MD07G1234700* (*WRKY70*) was localized directly within Module 19, consistent with this interspecies module’s strong functional enrichment for hypersensitive response and programmed cell death pathways. Crucially, signatures of positive selection for both *MD07G1234700* (*WRKY70*) and *MD01G1084700* (*DAG1*) were detected exclusively in *M. domestica* dessert varieties, with no signal of positive selection observed in cider varieties or wild populations.

**Fig. 6.**
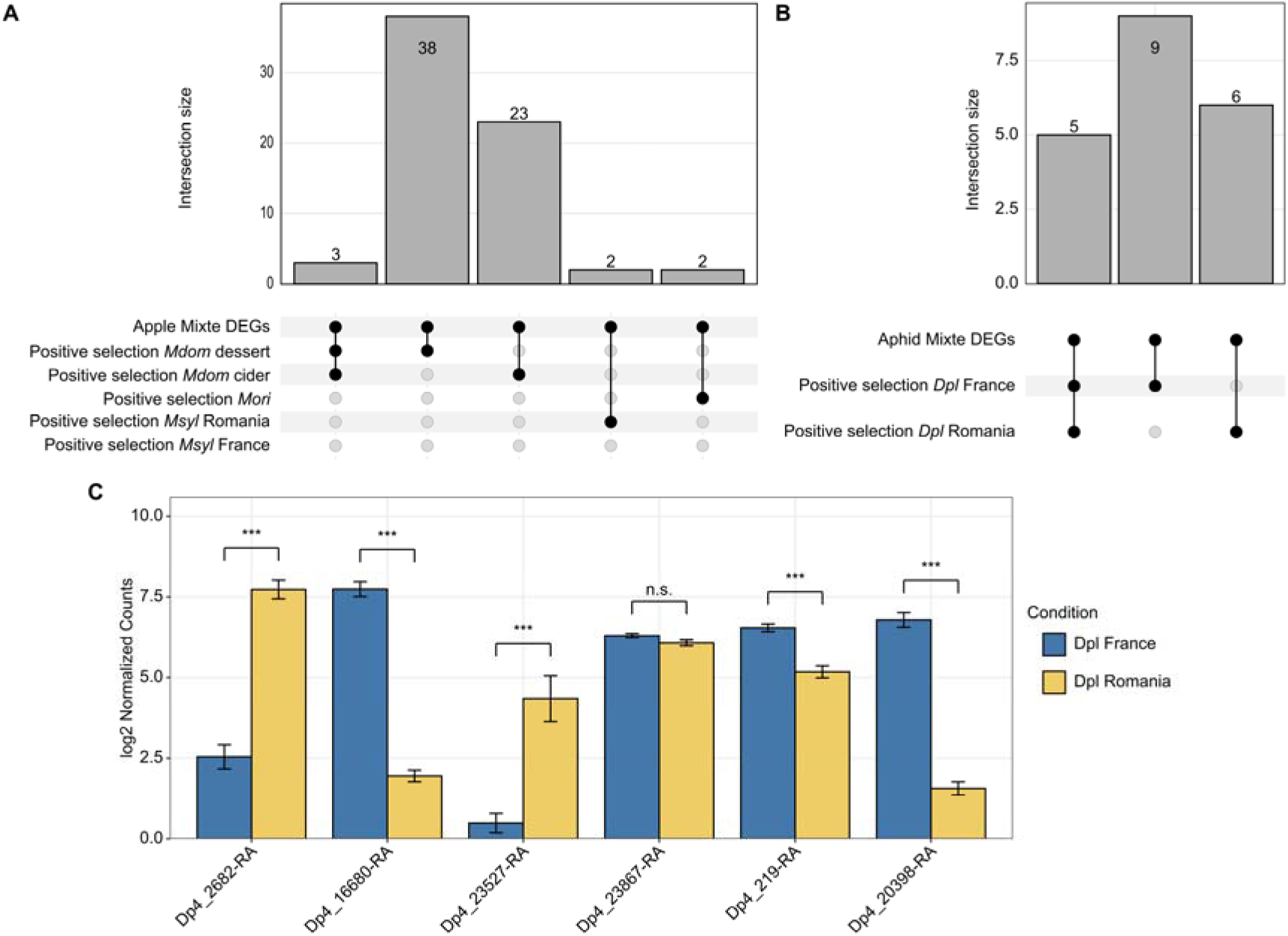
Positive selection targets aphid genes embedded in host-aphid interaction modules. (A) UpSet plot showing overlap between genes under positive selection and genes embedded within mixed host-aphid co-expression modules. (B) Distribution of population-specific positive selection signatures among aphid genotypes. Several candidate interaction genes exhibited lineage-specific selection patterns. (C) Expression profiles of candidate aphid genes associated with host interaction that are embedded in mixed modules and overlap positive-selection outlier regions. Heights represent mean log_2_ normalized counts (± SE) for Dpl France (blue) and Dpl Romania (yellow). Asterisks and labels above brackets denote statistically significant differences in expression between aphid genotypes based on false discovery rate thresholds (n.s. not significant; * FDR < 0.05; ** FDR < 0.01; *** FDR < 0.001). Alt Text: Three-panel figure illustrating overlaps between positive selection signatures and host-aphid interaction modules, alongside lineage-specific expression profiles for candidate aphid genes. Panel A and Panel B present UpSet plots detailing overlaps between genes in mixed co-expression modules and positive selection targets for apple populations and aphid genotypes, respectively. Panel C displays bar charts of log_2_ normalized counts comparing candidate aphid gene expression between Dpl France and Dpl Romania conditions Statistical significance for expression differences between aphid genotypes is indicated above the bars (n.s.: not significant; *: FDR < 0.05; **: FDR < 0.01; ***: FDR < 0.001).

### Positive selection targets aphid genes associated with host interaction

To investigate whether adaptive evolutionary pressures acted on the parasite side of these interspecies interaction networks, we evaluated selection signatures across aphid genes embedded within mixed host-aphid co-expression modules. Unlike host transcripts, we found no overlap between aphid genes with balancing-selection signatures and those assigned to mixed interspecies modules. Instead, we identified candidate positive-selection outliers among aphid genes in mixed modules. Across all mixed modules, 20 aphid genes displayed overlapping candidate positive-selection outlier regions in at least one aphid genotype (Fig. 6B) (Table S11).

Functional annotation using BLAST searches against dedicated receptor databases revealed that six of these 20 positively selected aphid genes shared significant homology with gustatory and/or salivary proteins (Table S11). None of the six candidate genes overlapped positive-selection outlier regions in both aphid populations. Specifically, positive selection targeted *Dp4_2682*, *Dp4_16680*, and *Dp4_23527* exclusively in the French aphid genotype, whereas *Dp4_23867*, *Dp4_219*, and *Dp4_20398* were positively selected solely in the Romanian population. Beyond sequence-level divergence, five of the six candidate receptor genes showed significant differences in transcript expression between French and Romanian aphid genotypes during host exploitation (Fig. 6C), with *Dp4_23867* the only gene not differentially expressed between the two genotypes. This pattern generally links lineage-specific directional selection to transcriptomic divergence.

Together, these results reveal population-specific associations between sequence-level selection signatures and expression divergence among aphid genes embedded in host–aphid co-expression modules.

## Discussion

### Altered aphid compatibility and host resistance between cultivated and wild apples

Our results reveal marked differences in aphid compatibility and resistance between wild and cultivated apples. Cultivated apples generally supported higher aphid performance, whereas Romanian *M. sylvestris* exhibited the strongest resistance, and each aphid genotype performed best on host populations, showing patterns partially consistent with shared recent evolutionary history. Importantly, aphid performance depended on the specific host-aphid genotype combination, revealing asymmetric host compatibility. The French aphid genotype performed best on cultivated apples, while the Romanian aphid genotype achieved higher fitness on wild populations, particularly those from Romania. Recent demographic reconstructions suggest that European aphid populations represent the ancestral lineage, followed by divergence in Mediterranean populations and subsequent expansion associated with the diffusion of cultivated apple. The spread of domesticated apples across Eurasia may therefore have created novel ecological opportunities promoting aphid specialization and divergence. Together, these results support the idea that domestication not only reshaped host defense strategies but also altered the selective landscape experienced by aphid populations. This is consistent with the broader view that crop domestication can restructure trophic interactions and drive adaptive changes in herbivore populations (Chen *et al*., 2015). These patterns align with host-population specialization following apple diversification, though we emphasize that our experimental design does not formally test host-aphid co-divergence. We note that formally testing host-aphid co-divergence would require a joint phylogenetic framework and broader sampling of host and aphid populations.

These findings align with broader evidence that domestication often reduces resistance to herbivores, although the magnitude and direction of these effects depend on the crop, the plant organ, and the herbivore guild (Turcotte *et al*., 2014; Whitehead *et al*., 2017). In particular, meta-analyses across numerous domestication events have shown that artificial selection for yield and productivity can come at the expense of defense, leading to increased susceptibility in many systems (Whitehead *et al*., 2017; Fernandez *et al*., 2021). However, the consequences of domestication for resistance are not uniform; some crops maintain or even enhance particular defense traits, especially against specialist herbivores that have coevolved with their hosts (Whitehead *et al*., 2017).

### Wild populations show patterns consistent with greater constitutive defense, and cultivated apples mount extensive inducible defense responses

Romanian *M. sylvestris* consistently exhibited the strongest resistance to aphids despite limited transcriptomic remodeling following infestation. This pattern suggests that resistance in wild populations may involve constitutive barriers or more efficient early responses that reduce the need for massive downstream transcriptional activation. The elevated flavonol levels observed in wild populations further support this hypothesis. Flavonoids are well-known secondary metabolites involved in stress responses and defense against herbivores, and their constitutively higher accumulation may contribute to reduced aphid performance. These results are consistent with the idea that domestication can alter both constitutive and inducible defense programs rather than simply weakening all defenses uniformly (Moreira *et al*., 2018). Indeed, previous work has shown that domesticated plants often exhibit reduced levels of both baseline and induced chemical defenses, making them more susceptible to herbivores (Moreira *et al*., 2018).

Our gene set overlap analyses further support potential differences in baseline defense regulation. Cross-referencing baseline DEGs with the core set of 1,201 *M. domestica*-specific infestation-responsive DEGs revealed significant enrichment in cultivated apples relative to *M. sylvestris* Romania and *M. orientalis*. Positive Z-scores across enriched biological processes suggest that pathways recruited upon infestation in cultivated varieties, particularly secondary metabolism and pathogen response, show higher baseline expression in certain wild populations under control conditions. This pattern is consistent with greater constitutive investment in these pathways. However, this pattern was not uniform across all comparisons: no enrichment was detected relative to French *M. sylvestris*, and functional enrichment was absent for dessert apples compared with *M. orientalis*. This subtle variation suggests that baseline expression patterns likely differ among wild populations rather than reflecting a uniform trend across lineages.

In perennial systems, wild relatives often harbor stronger or more diverse resistance traits than their cultivated counterparts, as documented for fire blight resistance in wild *Malus* genotypes (Emeriewen *et al*., 2017; Kostick *et al*., 2019). The retention of such variation in wild populations is facilitated by outcrossing, large effective population sizes, and recurrent gene flow among sympatric wild and domesticated gene pools (Cornille *et al*., 2013; Chen *et al*., 2026). Our system thus echoes a broader pattern in fruit trees, where wild relatives represent reservoirs of resistance alleles that have been partially eroded or reconfigured during domestication and breeding (Kumar *et al*., 2010; Bus *et al*., 2011). In our case, wild apples appear to rely on a defense strategy that minimizes transcriptional perturbation while effectively limiting aphid fitness, in contrast to cultivated varieties that mount extensive inducible responses yet remain more susceptible. An alternative, non-mutually exclusive explanation is that the muted transcriptional response in Romanian *M. sylvestris* reflects reduced inducibility rather than an adaptive constitutive defense strategy. Romanian *M. sylvestris* has previously been shown to carry an elevated load of deleterious mutations (Chen *et al*., 2026), raising the possibility that demographic processes have affected the efficiency of regulatory responses. Distinguishing between enhanced constitutive defense and reduced inducibility will require direct functional tests of defense signaling across wild genotypes.

An additional consideration is that cultivated apples were grafted onto M9 rootstocks, reflecting standard horticultural practice, whereas wild seedlings were grown on their own roots. Scion–rootstock interactions can influence plant physiology and gene expression, and some of the baseline transcriptomic divergence observed between cultivated and wild apples may therefore reflect both genetic changes associated with domestication and cultivation-related effects of grafting. Because grafting is integral to modern apple production, this contrast can be viewed as part of the broader cultivated phenotype examined here, although the present design cannot disentangle the respective contributions of scion genotype and rootstock effects. Together, the divergence in baseline expression suggests that cultivated and wild apples differ not only in infestation-induced responses but also in regulatory programs expressed before attack, potentially reflecting the combined effects of domestication history and cultivation practices.

The extensive transcriptional response of cultivated apples provides the complementary side of this contrast. Despite mounting thousands of infestation-responsive changes, cultivated apples supported greater aphid abundance than wild populations. This suggests that response magnitude does not necessarily translate into effective resistance and that domestication may have altered the balance between baseline and inducible defense investment. Our findings align with the broader pattern that domestication can reduce the efficiency or specificity of inducible defenses, potentially because relaxed herbivore selection in managed environments or trade-offs favoring growth and yield (Whitehead *et al*., 2017; Moreira *et al*., 2018). Cider varieties also showed repression of photosynthesis-associated pathways during infestation, consistent with, although not sufficient to demonstrate, resource reallocation associated with defense activation.

Interestingly, recent pan-genomic analyses in apple have revealed that cultivated genotypes carry more resistance gene analogs (RGAs) than their wild relatives, owing in part to segmental and tandem duplications and to introgression from multiple wild species (Su *et al*., 2024; Chen *et al*., 2026). This suggests that, despite an overall increase in susceptibility to aphids, the cultivated apple genome has retained or even expanded certain components of the immune repertoire, possibly in response to other pathogens such as *Venturia inaequalis* or *Erwinia amylovora* (Bus *et al*., 2011; Kostick *et al*., 2019). Overall, these results demonstrate that perennial crop domestication can generate distinct defense syndromes, with cultivated apples relying on large-scale inducible responses that may be less effective at constraining aphid performance, even as other arms of the immune system are maintained or expanded.

### Cross-species co-expression modules link host defense-signalling genes to candidate aphid effectors

Integrated co-expression analyses identified mixed host-aphid regulatory modules that may underlie coevolutionary interactions. Module 19 represents a particularly strong candidate interaction hub because it combines defense-associated host genes, candidate aphid effectors, signatures of balancing selection, and enrichment for hypersensitive response genes. The coexistence of host resistance genes and aphid-derived genes within the same module suggests correlated cross-species expression patterns. Balancing selection detected on several host resistance genes further suggests long-term maintenance of polymorphism, potentially driven by fluctuating parasite-mediated selection. These findings echo theoretical expectations that host-parasite coevolution can maintain genetic diversity at resistance loci through negative frequency-dependent selection or spatially and temporally variable selection pressures (Tellier and Brown, 2007; Penczykowski *et al*., 2015).

The positive-selection analyses provide a complementary signal on the host side. Several infestation-responsive genes overlapping candidate positive-selection regions were specific to cultivated apples, including homologs of the transcriptional regulators WRKY70 and DAG1. WRKY70 is particularly notable because it was embedded within Module 19 and is involved in salicylic acid–jasmonic acid crosstalk and defense regulation in *Arabidopsis* (Besseau *et al*., 2012). This convergence between inducible expression, module membership, and lineage-specific selection suggests that domestication-associated evolution may have targeted upstream regulatory components of defense rather than simply reducing resistance gene content.

On the aphid side, candidate genes overlapping positive-selection outliers showed lineage-specific patterns, with different sets of gustatory and salivary-associated genes implicated in French and Romanian aphids. Five of the six candidate genes also differed in expression between aphid lineages. Although these associations do not establish direct host-driven selection, they are consistent with geographically distinct selective environments acting on sensory and salivary functions involved in host exploitation. Such divergence may have been reinforced during the expansion of cultivated apple and the subsequent differentiation of aphid populations.

In perennial crops, long generation times, clonal propagation, and recurrent introgression from wild relatives are expected to shape the dynamics of resistance alleles in distinctive ways (Miller and Gross, 2011; Gaut *et al*., 2015). For instance, genome-wide scans in apples have revealed that dessert cultivars exhibit more hard selective sweeps at disease resistance and flowering-time genes, whereas cider apples show proportionally more soft sweeps and signatures of balancing selection (Chen *et al*., 2026). Such patterns are consistent with multiple domestication routes and divergent selection pressures acting on different cultivated gene pools. Our results thus provide empirical support for the idea that domestication not only reshapes host defense strategies but also alters the selective landscape experienced by aphid populations, potentially driving the evolution of novel effector repertoires and sensory adaptations.

## Conclusion

Our study demonstrates that apple domestication shifted the balance between constitutive and inducible defense signaling. Wild populations exhibited greater resistance with weaker transcriptional perturbation, consistent with a more constitutive defense strategy, whereas cultivated apples mounted extensive inducible transcriptional reprogramming of hormone-signaling and secondary-metabolism pathways yet remained more susceptible. Cross-species co-expression analyses further identified candidate regulatory modules, including a WRKY70 homolog and a CNL-class resistance gene, linking host defense-signaling genes to aphid transcripts with effector-like annotations, offering a mechanistic entry point into these host-aphid interactions. By pairing wild and cultivated apple populations with aphid genotypes of matched geographic origin, our reciprocal design, rarely achieved in perennial crop-pest systems, further allowed us to test for, and identify candidate signatures of, local host-aphid coadaptation. Our findings in apple, a long-lived, clonally propagated perennial fruit tree, thus provide a test case for how domestication reshapes induced immunity to phloem-feeding insects in crops with complex evolutionary histories and extensive gene flow from wild relatives, with implications for future breeding and conservation programs.

## Supporting information

Supplementary informations

## Supplementary Data

The following Supplementary Data are available for this article and are referenced in the main text.

Table S1. Details of seedlings, including origin, condition, position during the experiment, and information collected during the experiment.

Table S2. Details of apple RNA-seq sample filtering and mapping statistics.

Table S3. Model results for leaf phenotypic traits and aphid counts at sampling time.

Table S4. Pairwise contrasts of leaf phenotypic traits per level of condition, separated by apple population.

Table S5. Aphid count analyses, including host population and genotype pairwise contrasts, the host domestication status GLMM summary, and domestication-status estimated marginal means and contrasts.

Table S6. Gene set overlap between core inducible defense DEGs and baseline transcriptomic divergence across apple populations.

Table S7. Selection signals and functional annotations for co-expressed genes in Host-effect Cluster 19.

Table S8. Selection signals and functional annotations for co-expressed genes in Host-effect Cluster 49.

Table S9. Table of apple genes present in mixed co-expression clusters with balancing selection signals and their annotation.

Table S10. Table of apple genes present in mixed co-expression clusters with positive selection signals and their annotation.

Table S11. Table of aphid genes present in mixed co-expression clusters with positive selection signals and their annotation.

Figure S1. Visual classification categories for aphid identification.

Figure S2. DHARMa dispersion test for the aphid count Poisson GLMM.

Figure S3. Partitioning of total gene expression variance across experimental factors.

Figure S4. Domestication altered host physiological responses: leaf chlorophyll content.

Figure S5. Domestication altered host physiological responses: Nitrogen Balance Index (NBI).

Figure S6. Expression profile of AGAMOUS-LIKE 42 (AGL-42) homologs across cultivated and wild *Malus* populations.

Figure S7. Gene Ontology (GO) enrichment of *M. domestica*-specific inducible genes differentially expressed at baseline relative to wild species.

Figure S8. Bubble plot summarizing enriched biological processes associated with DEGs in response to infestation specific to the *M. domestica* cider variety.

## Acknowledgements

This research was funded by the ATIP-CNRS, Inserm, IDEEV, and LabEx BASC; by Tamkeen, under grant number AD454 from the New York University Abu Dhabi Research Institute; and ANR-21-CE20-0005 (PLEASURE), led by A.C. RD was supported by a PhD fellowship from the French Ministry of Higher Education and Research through the Graduate School ED SEVE 567 (Plant Sciences: From Genes to Ecosystems). The authors thank the INRAE Biological Resource Center “Pome Fruits and Roses” (RosePom) and associated staff, especially the INRAE Horticulture Experimental Facility (Unité Expérimentale Horticole, INRAE, 49000 Angers, France), for plant material and technical support. We also thank Frederique Didelot, Arnaud Lemarquand, Christian Cattanéo and Remi Gardet (PHENOTIC, INRAE Angers, 42 rue Georges Morel, 49070 Beaucouzé, France) for phenotyping platform support. AR and TU received funding through project number 88-PHE (PN-IV-P8-8.1-PRE-HE-ORG-2024-0223) and RECOVER (PN-IV-P6-6.1-CoEx-2024-0124) from the MCDI.

This research was carried out on the High-Performance Computing resources at New York University Abu Dhabi and on the Core Cluster of the Institut Français de Bioinformatique (IFB) (ANR-11-INBS-0013). We thank Adrien Falce, Olivier Langella, and Benoit Johannet for their help and support at the INRAE Génétique Quantitative et Evolution - Le Moulon lab.

## Competing interests

The authors declare no competing interests.

## Author contributions

AC designed the experiments and acquired funding. AH, AM, AN, AR, TU and AC collected materials and provided plant and aphid genetic resources. XC, AV, AM, AH, SO-V, JY and AC performed plant phenotyping and aphid infestation experiments. AV, JY and AC prepared samples and conducted RNA sequencing. RD performed statistical modelling of phenotypic and aphid count data. RD and XC analyzed transcriptomics, co-expression networks, and selection scans. RD, MC, XC, NCeS and AC interpreted the results. RD and AC wrote the original draft. RD, XC, NCeS, and AC reviewed and edited the manuscript, with contributions from all authors.

## Data availability

Host genomic variant data and baseline selection signatures used in this study were obtained from Chen *et al*., 2026, deposited under NCBI BioProject PRJNA1252109. RNA-seq data generated for the present infestation experiment (host and aphid) have been deposited in NCBI under BioProject accession PRJNA1534890. Phenotypic and read count data used to run analyses are available on Zenodo 10.5281/zenodo.22230915. The analysis code and workflow are available at https://github.com/CornilleEclecticLab/Apple-RNAseq_EXP2. The lead contact can provide any additional information needed to reanalyze the data reported in this paper.

## Notes

### Competing Interest Statement

The authors have declared no competing interest.

### Summary of Updates

- Change to manuscript formatting - Addition of more precision to some figure legends - Change of figure panel convention from (a-b) to (A-B) - Removal of individuals linked to old experimental planning from Table S1 who were not used in analysis.

