## Supplementary informations for "Domestication reshaped induced defense signaling in the apple-aphid interaction"

### *Journal of Experimental Botany* Supplementary Data

Article acceptance date: Click here to enter a date.

The following Supporting Information is available for this article:

**Fig. S1.** Visual classification categories for aphid identification.

**Fig. S2.** DHARMa dispersion test for the aphid count Poisson GLMM.

**Fig. S3.** Partitioning of total gene expression variance across experimental factors.

**Fig. S4.** Domestication altered host physiological responses: leaf chlorophyll content.

**Fig. S5**. Domestication altered host physiological responses: Nitrogen Balance Index (NBI).

**Fig. S6.** Expression profile of AGAMOUS-LIKE 42 (AGL-42) homologs across cultivated and wild *Malus* populations.

**Fig. S7.** Gene Ontology (GO) enrichment of *M. domestica*-specific inducible genes differentially expressed at baseline relative to wild species.

**Fig. S8.** Bubble plot summarizing enriched biological processes associated with DEG in response to infestation specific to the *M. domestica* cider variety.

**Table S1.** Details of seedlings with origins, condition, position during experiment, and information collected during experiment.

**Table S2.** Details of apple RNA-seq sample filtering and mapping statistics.

**Table S3.** Model results for leaf phenotypic traits and aphid counts at sampling time.

**Table S4.** Pairwise contrasts of leaf phenotypic traits per level of condition, separated by apple population.

**Table S5.** Aphid count analyses including host population and genotype pairwise contrasts, host domestication status GLMM summary, and domestication status estimated marginal means and contrasts.

**Table S6.** Gene set overlap between core inducible defense DEGs and baseline transcriptomic divergence across apple populations.

**Table S7.** Selection signals and functional annotations for co-expressed genes in Host-effect Cluster 19.

**Table S8.** Selection signals and functional annotations for co-expressed genes in Host-effect Cluster 49.

**Table S9.** Table of apple genes present in mixed co-expression clusters with balancing selection signals and their annotation.

**Table S10.** Table of apple genes present in mixed co-expression clusters with positive selection signals and their annotation.

**Table S11.** Table of aphid genes present in mixed co-expression clusters with positive selection signals and their annotation.

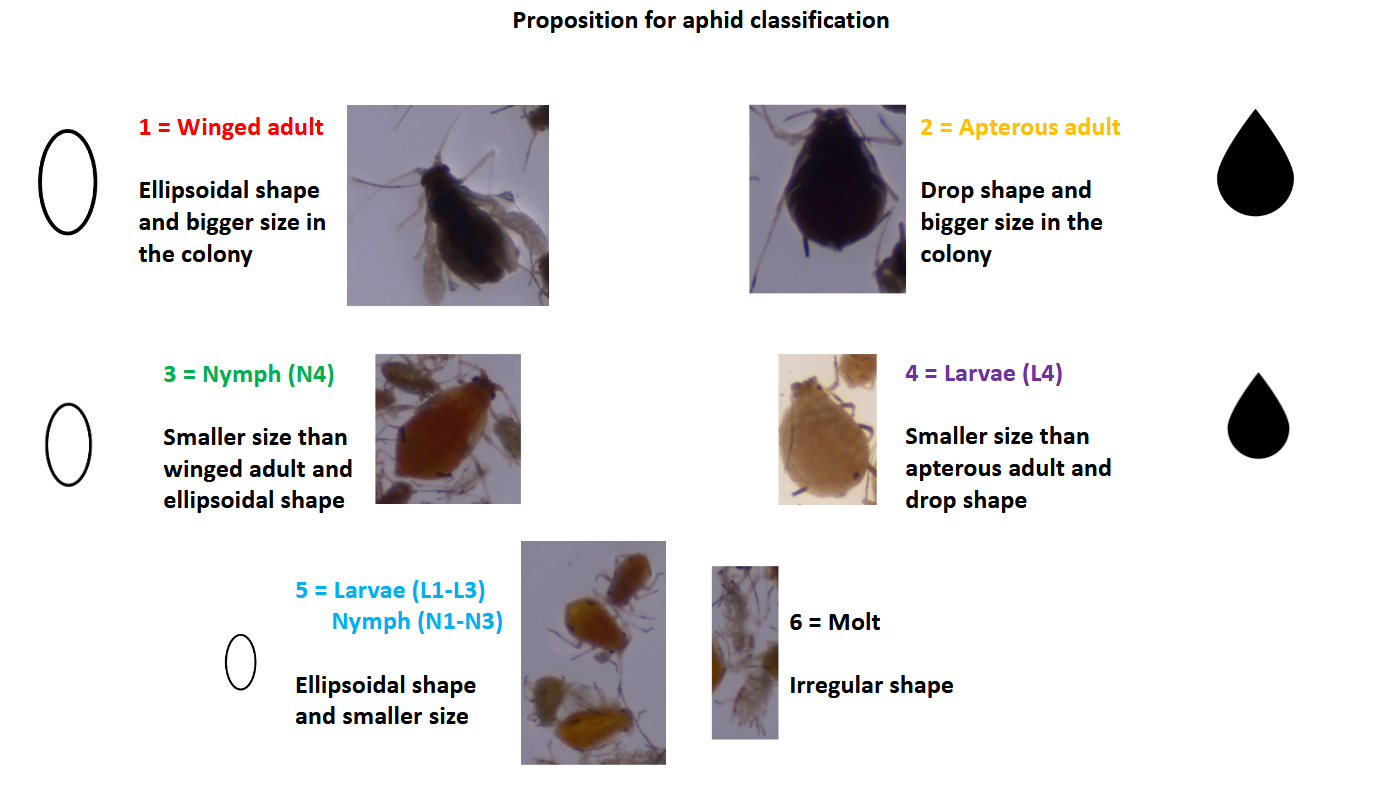

**Figure S1. Visual classification categories for aphid identification.** Aphid categorization scheme used for visual scoring, showing the six defined classes: (1) winged adult, (2) apterous adult, (3) nymph precursor of winged form, (4) larva precursor of apterous form, (5) larva, and (6) molt. Adapted from Olvera Vázquez (2023); originally developed with expert input from Dr. Marcos Miñarro (SERIDA) and Dr. Ahmar Ahlmedi (pcfruit).

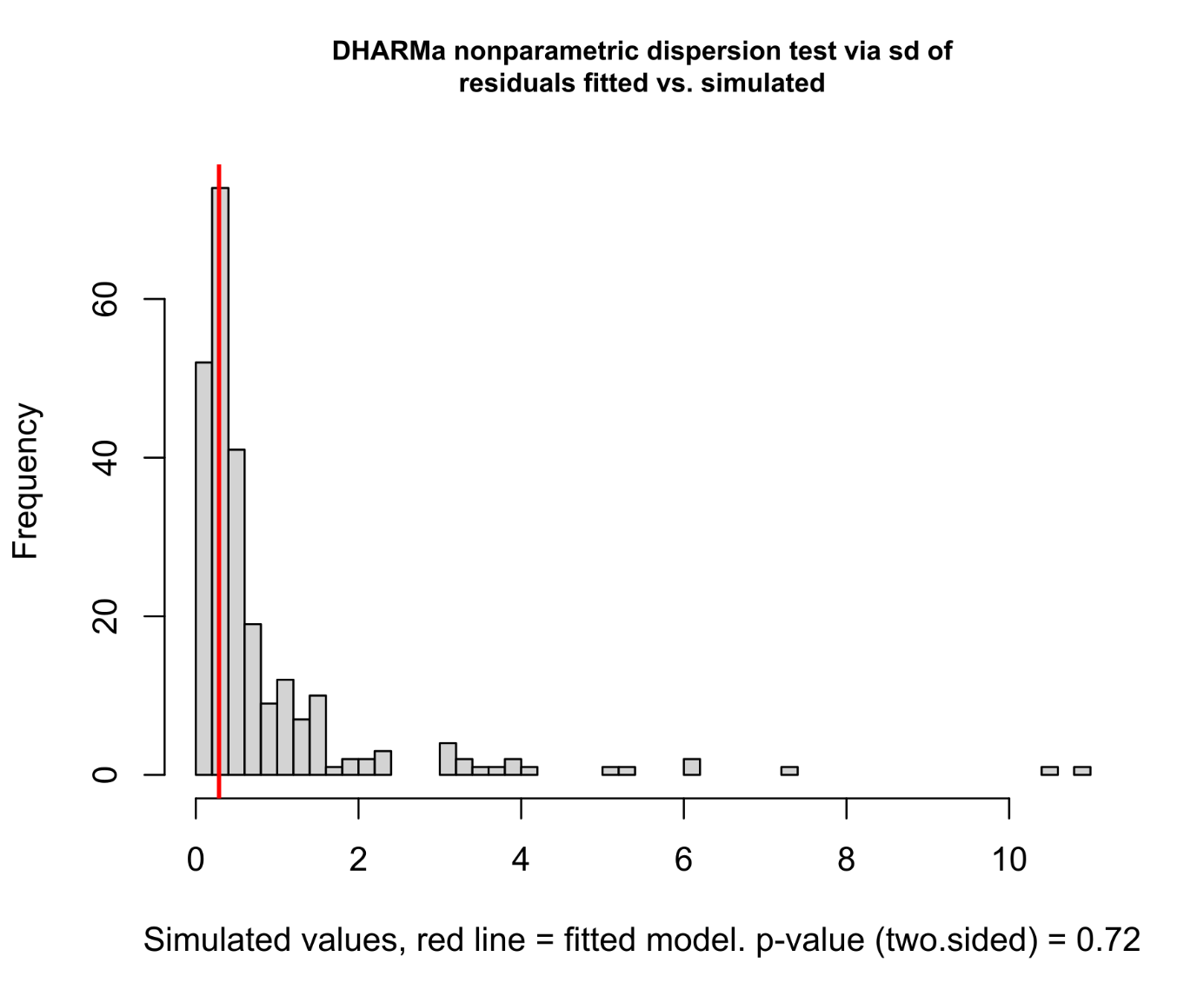

**Figure S2. DHARMa dispersion test for the aphid count Poisson GLMM.** Non-parametric dispersion test output generated via DHARMa::testDispersion() evaluating residual variance in the model predicting aphid counts. The histogram shows the distribution of expected dispersion statistics calculated across simulations (N = 250). The vertical red line marks the observed dispersion statistic (Dispersion ratio 0.33589, p = 0.72). The non-significant p-value (p > 0.05) indicates no evidence of overdispersion or underdispersion, confirming the appropriateness of the Poisson error distribution.

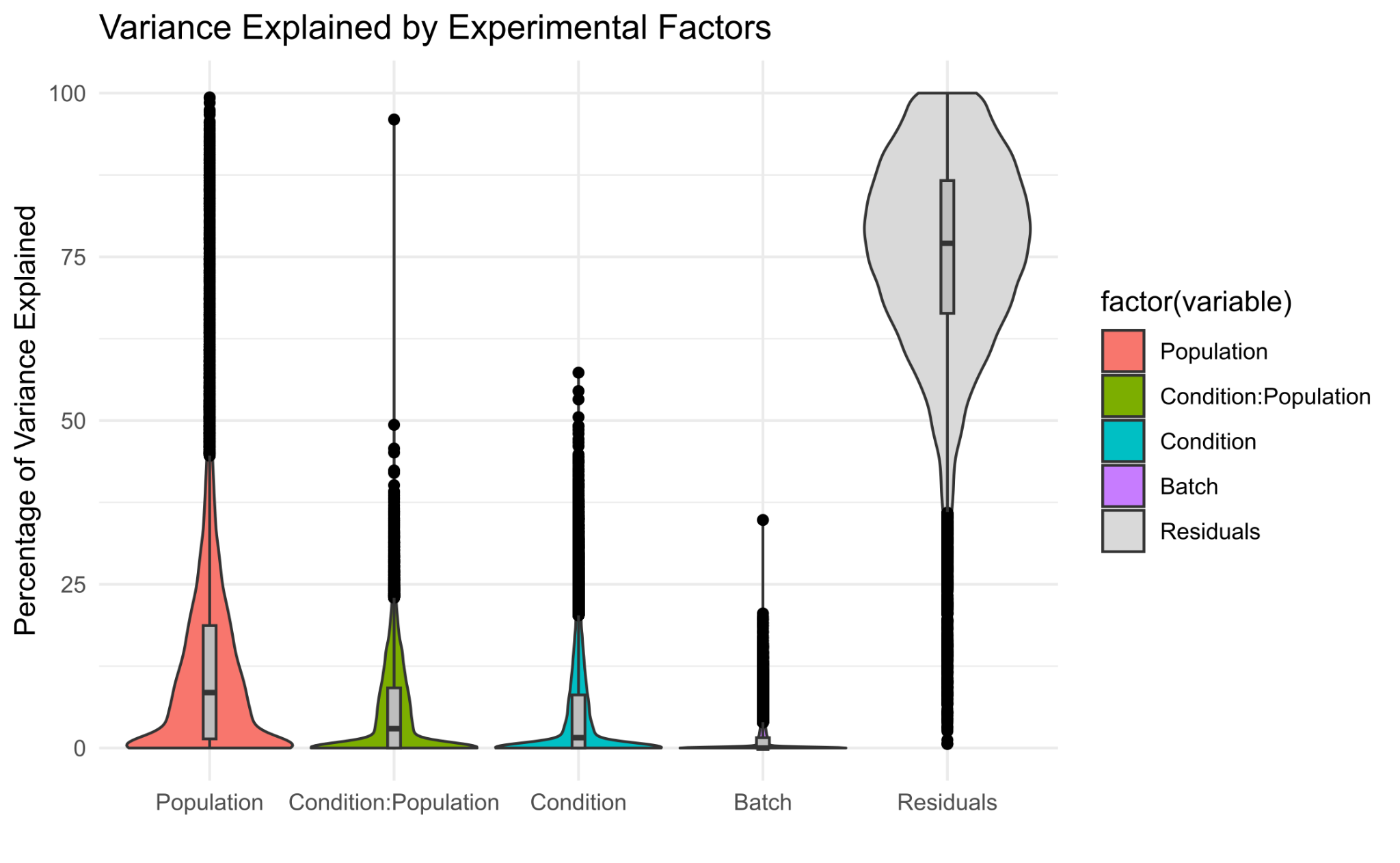

**Figure S3. Partitioning of total gene expression variance across experimental factors.** Violin plots illustrating the distribution of variance explained by biological and technical covariates across 24,413 filtered genomic features, estimated using a linear mixed-effects model in the variancePartition R package. Evaluated factors include experimental infestation condition (Condition), apple seedling population origin (Population), their interaction (Condition:Population), experimental batch (Batch), and unexplained residual variation (Residuals). Within each violin, boxplots display the median (horizontal line) and interquartile range of variance explained across all analysed genes. Technical batch effect accounted for a negligible proportion of global transcriptomic variation (mean = 1.20%, median = 0.00%), validating that sample processing across two batches did not introduce systematic bias or confound downstream differential expression analyses.

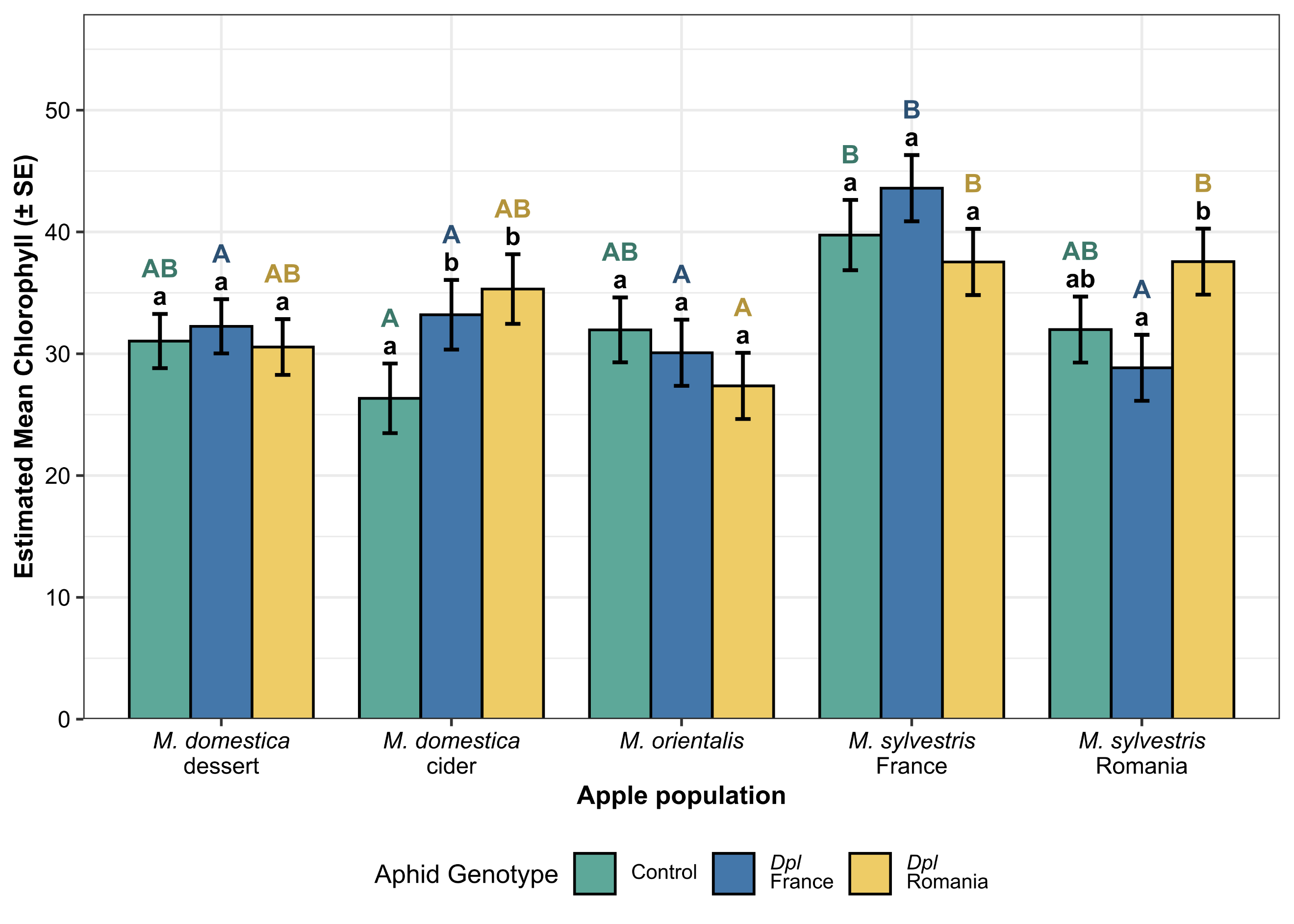

**Figure S4. Domestication altered host physiological responses: leaf chlorophyll content.** Estimated mean leaf chlorophyll content (Chl) (± SE) across five apple populations (*Malus domestica*, *M. orientalis*, and *M. sylvestris*) subjected to three conditions: uninfested control (green), infested by French aphid genotype (blue), and infested by Romanian aphid genotype (yellow). Bar heights represent the estimated marginal means (emmeans) derived from the interaction models. Error bars represent the standard error of the mean. Different letters above the bars indicate statistically significant differences based on Tukey’s HSD post-hoc tests (p < 0.05). Black lowercase letters denote significant differences between the aphid treatment conditions within a single given apple population. Colored uppercase letters denote significant differences across the five apple populations within a specific aphid treatment condition. Complete statistical results are provided in Tables S3–S4.

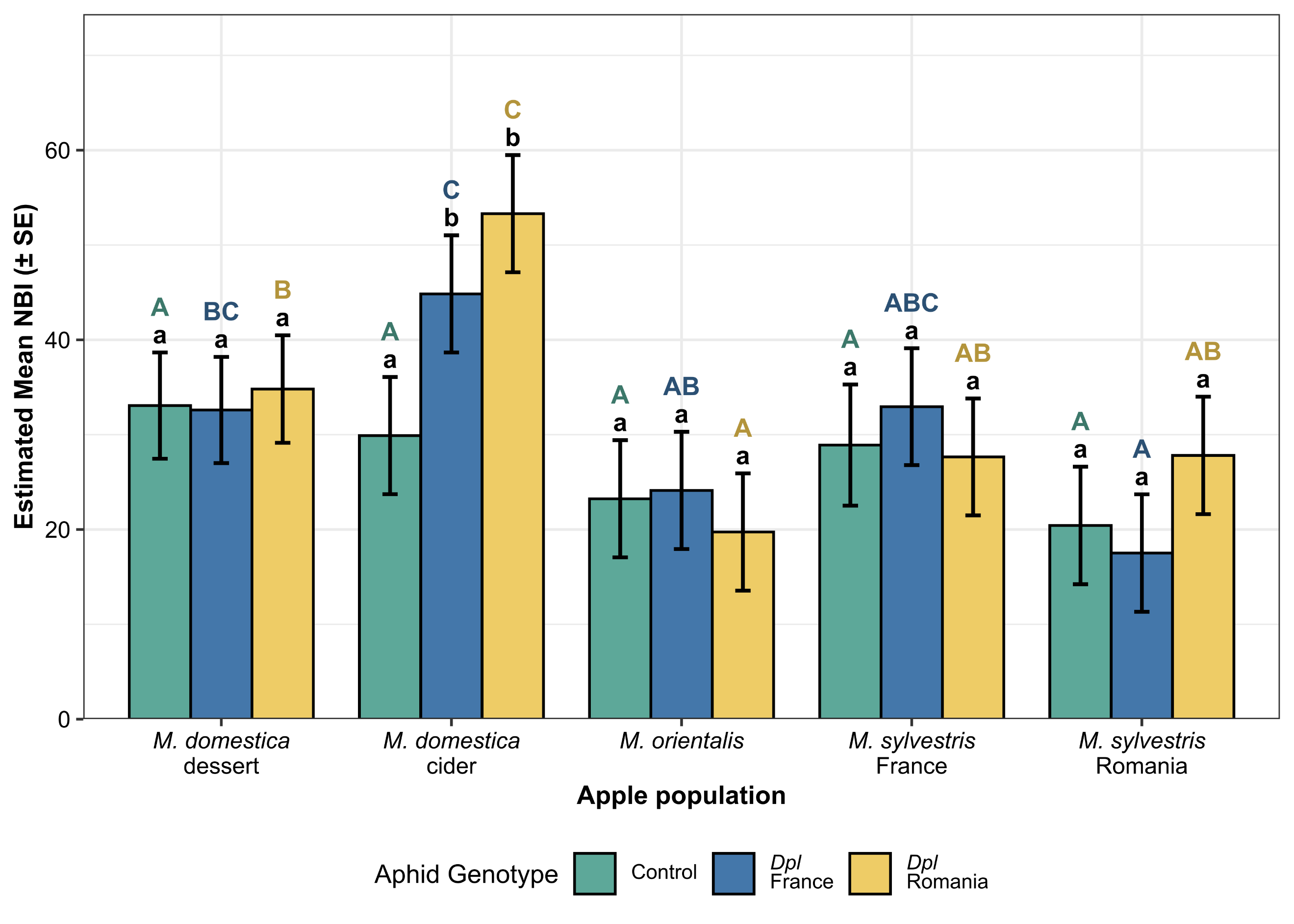

**Figure S5. Domestication altered host physiological responses: Nitrogen Balance Index (NBI).** Estimated mean Nitrogen Balance Index (NBI) (± SE) across five apple populations (*Malus domestica*, *M. orientalis*, and *M. sylvestris*) subjected to three conditions: uninfested control (green), infested by French aphid genotype (blue), and infested by Romanian aphid genotype (yellow). Bar heights represent the estimated marginal means (emmeans) derived from the interaction models. Error bars represent the standard error of the mean. Different letters above the bars indicate statistically significant differences based on Tukey’s HSD post-hoc tests (p < 0.05). Black lowercase letters denote significant differences between the aphid treatment conditions within a single given apple population. Colored uppercase letters denote significant differences across the five apple populations within a specific aphid treatment condition. Complete statistical results are provided in Tables S3–S4.

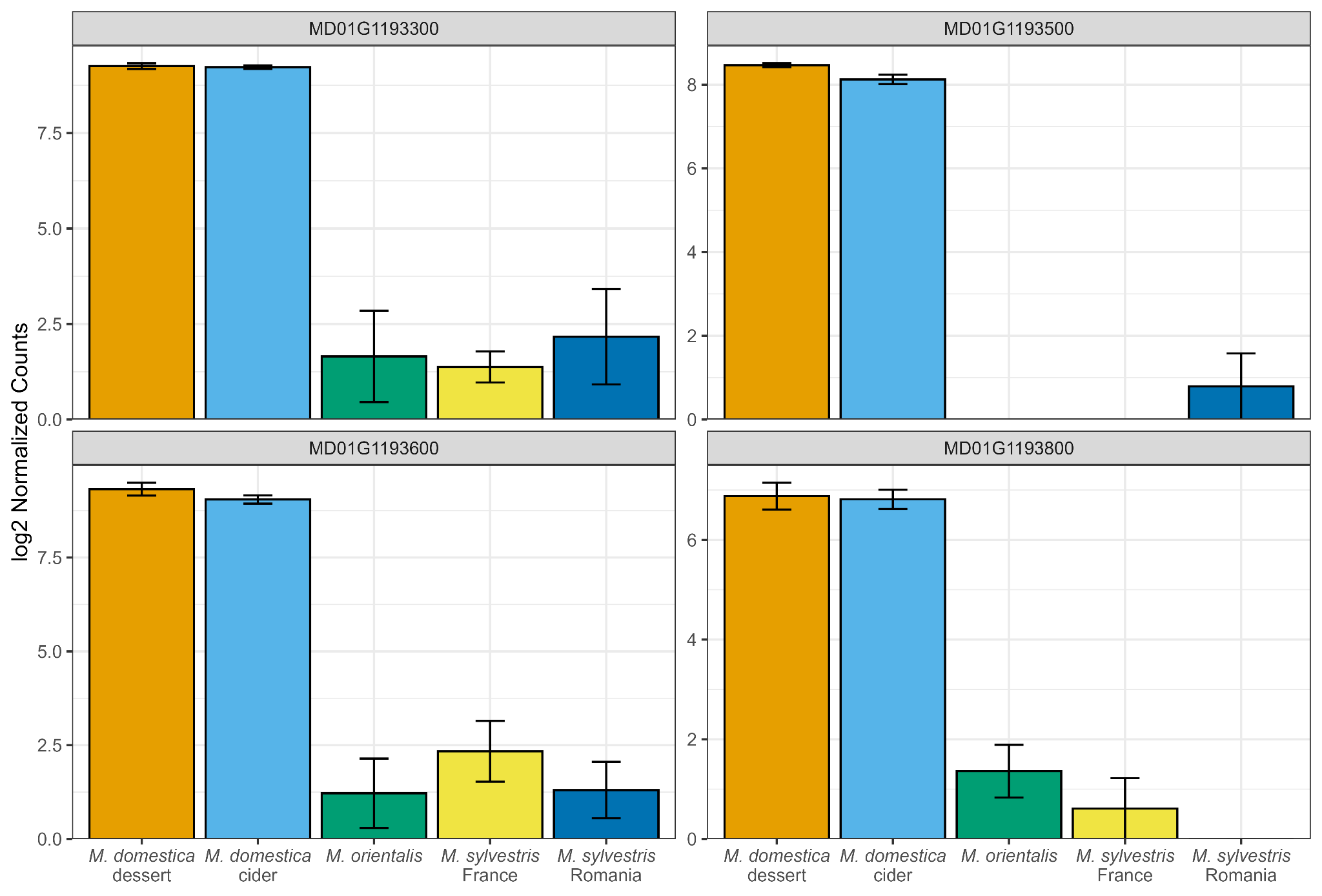

**Figure S6. Expression profile of *AGAMOUS-LIKE 42* (*AGL-42*) homologues across cultivated and wild *Malus* populations.** Bar plots represent mean log_2-normalised counts (+-SE) for four *Malus domestica*-specific differentially expressed genes (DEGs) annotated as *AGL-42* (MD01G1193300, MD01G1193500, MD01G1193600, and MD01G1193800) under control conditions.

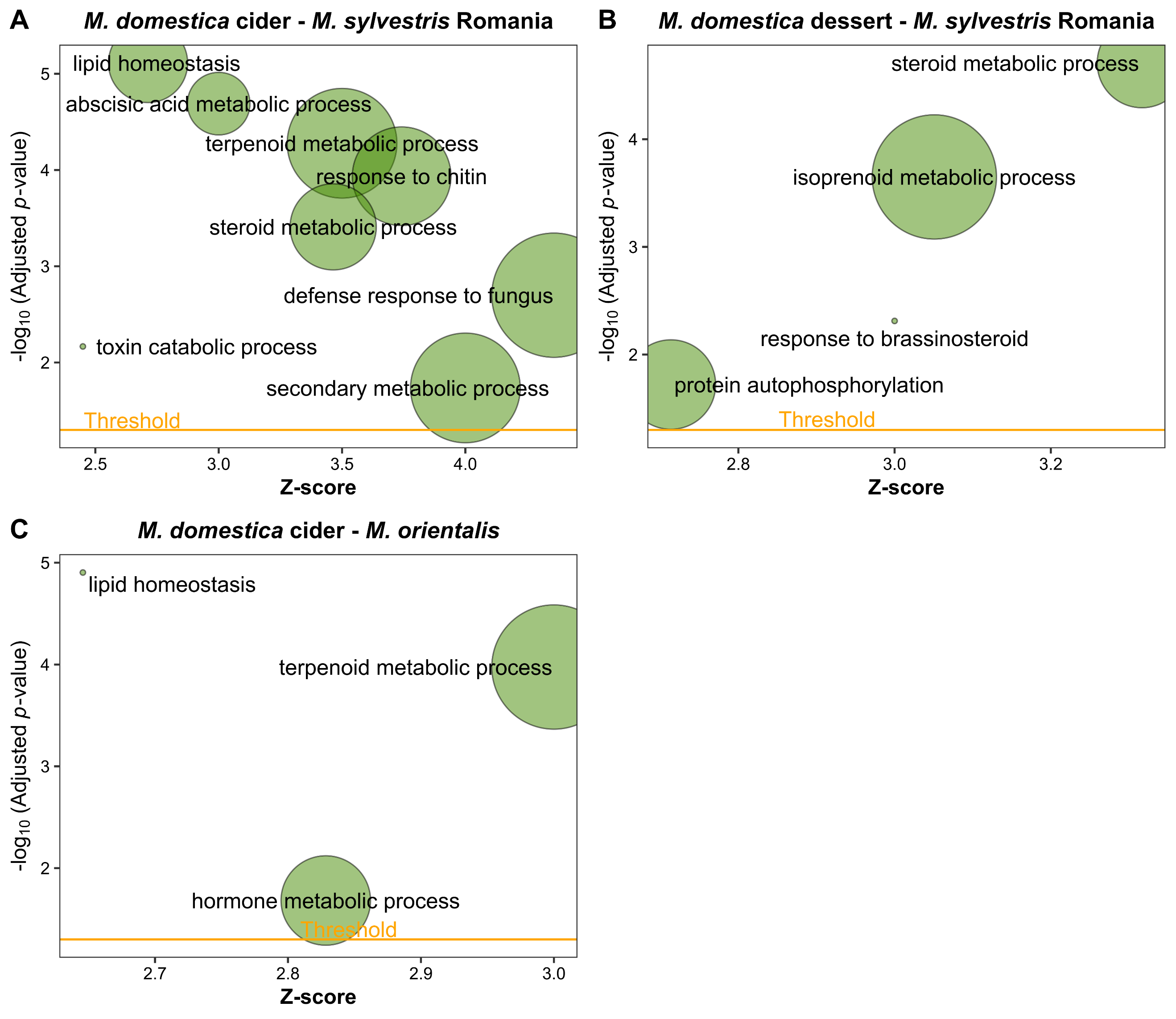

**Figure S7. Gene Ontology (GO) enrichment of *M. domestica*-specific inducible genes differentially expressed at baseline relative to wild species.** Bubble plots display enriched Biological Process (BP) terms (p-adj < 0.05) for: (A) *M. domestica* cider vs. *M. sylvestris* Romania, (B) *M. domestica* dessert vs. *M. sylvestris* Romania, and (C) *M. domestica* cider vs. *M. orientalis*. The y-axis indicates significance (-log_10_ adjusted p-value), the x-axis represents the Z-score (where positive Z-scores denote higher baseline expression in wild species relative to *M. domestica*), and bubble size is proportional to the number of assigned genes. The orange horizontal line marks the significance threshold (p-adj = 0.05).

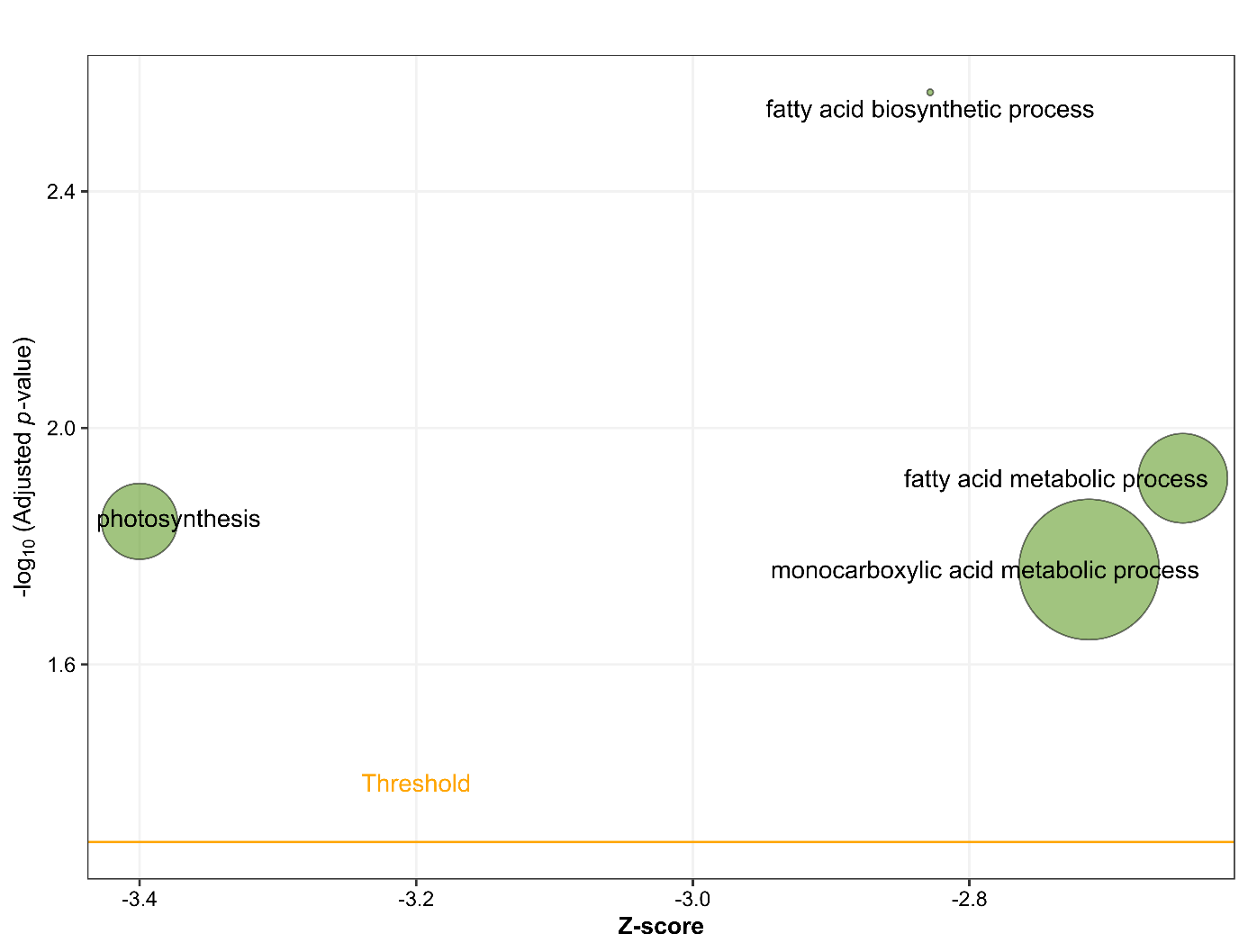

**Figure S8. Bubble plot summarising enriched biological processes associated with DEG in response to infestation specific to the *M. domestica* cider variety.** The y-axis represents statistical significance measured as log_10_(adjusted p-value). The x-axis indicates the GO term z-score. Bubble size is proportional to the total number of genes mapped to the GO term.
